# On-demand low-frequency stimulation for seizure control: efficacy and behavioral implications

**DOI:** 10.1101/2023.04.17.537172

**Authors:** Enya Paschen, Piret Kleis, Diego M. Vieira, Katharina Heining, Christian Boehler, Ulrich Egert, Ute Häussler, Carola A. Haas

**Affiliations:** Experimental Epilepsy Research, Department of Neurosurgery, Medical Center - University of Freiburg, Faculty of Medicine, Freiburg, Germany; Faculty of Biology, University of Freiburg, Freiburg, Germany; Biomicrotechnology, Department of Microsystems Engineering – IMTEK, Faculty of Engineering, University of Freiburg, Freiburg, Germany; Department of Neuroscience, Karolinska Institutet, Stockholm, Sweden; Department of Microsystems Engineering (IMTEK), Bioelectronic Microtechnology (BEMT), University of Freiburg, Freiburg, Germany; BrainLinks-BrainTools Center, University of Freiburg, Freiburg, Germany

**Keywords:** deep brain stimulation, kainate, hippocampus, closed-loop, learning and memory, spatial navigation

## Abstract

Mesial temporal lobe epilepsy (MTLE), the most common form of focal epilepsy in adults, is often refractory to medication and associated with hippocampal sclerosis. Deep brain stimulation represents an alternative treatment option for drug-resistant patients who are ineligible for resective brain surgery. In clinical practice, closed-loop stimulation at high frequencies is applied to interrupt ongoing seizures, yet with a high incidence of false detections, the drawback of delayed seizure-suppressive intervention and limited success in sclerotic tissue. More recently, hippocampal low-frequency stimulation (LFS) has been shown to reduce excitability in clinical settings and prevent seizures in experimental MTLE when applied continuously. However, as the hippocampus is important for navigation and memory, it would be beneficial to stimulate it only on-demand to reduce its exposure to LFS pulses, and to investigate LFS-related effects on cognition.

Using the intrahippocampal kainate mouse model, which recapitulates the key features of MTLE, we developed an on-demand LFS setup and investigated its effects on spontaneous seizure activity and hippocampal function. Specifically, our online detection algorithm monitored epileptiform activity in hippocampal local field potential recordings and identified short epileptiform bursts preceding focal seizure clusters, triggering hippocampal LFS to stabilize the network state. In addition, we investigated the acute influence of LFS on behavioral performance, including anxiety-like behavior in the light-dark box test, spatial and non-spatial memory in the object location memory and novel object recognition test, as well as spatial navigation and long-term memory in the Barnes maze.

Compared to open-loop stimulation protocols, on-demand LFS was more efficient in preventing focal seizure clusters, as the strong anti-epileptic effect was achieved with a reduced stimulation load. In behavioral tests, chronically epileptic mice were as mobile as healthy controls but showed increased anxiety, an altered spatial learning strategy and impaired memory performance. Most importantly, our experiments ruled out deleterious effects of hippocampal LFS on cognition and even showed alleviation of deficits in long-term memory recall. Taken together, our findings may provide a promising alternative to current therapies, overcoming some of their major limitations, and inspire further investigation of LFS for seizure control in MTLE.

## Introduction

Epilepsy is a prevalent chronic neurological disorder, affecting 0.5-1% of the world’s population ^1^. The most frequent form of focal epilepsies in adults is mesial temporal lobe epilepsy (MTLE) in which 30% of patients suffer from drug-resistant seizures ^2^ and cognitive comorbidities such as memory deficits ^3–5^. The major histopathological hallmark of MTLE is hippocampal sclerosis (HS), characterized by neuronal cell loss and reactive gliosis. HS is often associated with granule cell dispersion (GCD) and mossy fiber sprouting ^6, 7^. In many cases, surgical removal of the epileptogenic focus represents the only curative solution ^8, 9^. However, patients with multiple seizure foci or those at risk of resection-related impairments have limited treatment options, demonstrating an urgent need for new therapeutic approaches.

Neuromodulation via deep brain stimulation (DBS) provides new treatment avenues for conventionally untreatable patients. Electrical neuromodulation excels in terms of spatial and temporal precision compared to pharmacological intervention ^10^. Compared to resective surgery, DBS is reversible, adjustable, and potentially less invasive ^11, 12^. There are two main approaches, open-loop and closed-loop stimulation, that are used to control seizures ^13^. Open loop electrical high-frequency stimulation (HFS, 100–200 Hz) is delivered in a preprogrammed manner independent of ongoing brain activity or seizure occurrence (for review see ^14^). Closed loop systems, like responsive neurostimulation (RNS^®^), trigger stimulation once a seizure is detected ^15–17^, aiming to disrupt seizure propagation ^12^. Especially for MTLE patients with HS, HFS has low effectiveness ^14, 18, 19^, presumably due to extensive neuronal loss, reactive gliosis, and the resulting change in electrical resistance in the sclerotic hippocampus ^20, 21^. In contrast, hippocampal low-frequency stimulation (LFS) at 5 Hz reduced seizure activity remarkably in small clinical cohort studies, including MTLE patients with HS ^22–24^. Additionally, recent findings demonstrated that LFS (10 min on/off) at 1 Hz in the epileptic focus decreased cortical excitability in pharmacoresistant epilepsy patients ^25^. Due to the low intrinsic epileptogenic threshold of the hippocampal formation, LFS might be advantageous over HFS since it has a lower probability to elicit generalized seizures when applied in an interictal phase ^26^.

The medial temporal lobe, with its hippocampal-entorhinal circuitry, is the hub of learning, spatial navigation, and memory ^27, 28^, thus neuromodulation by DBS may affect these cognitive functions. In this regard, conflicting results were reported for HFS and RNS, likely due to variations in stimulation target, protocol, and the cognitive test applied (for review see ^10, 29, 30^). However, little is known about LFS-related cognitive effects ^22, 23^. For a systematic assessment of the LFS-related impact on seizure reduction and hippocampus-dependent cognitive functions, studies in translational animal models are crucial.

Several studies used open-loop LFS to interfere with seizures in chronically epileptic rodents ^31–33^. In fact, our previous work in intrahippocampal kainate (ihKA)-treated mice supports the hypothesis that the timing and frequency of hippocampal stimulation determine whether the effect is pro or antiepileptic ^34, 35^. The ihKA mouse model is well-accepted for the investigation of MTLE since it exhibits the major hallmarks of human pathology, comprising spontaneous recurrent epileptiform activity and histopathological changes in form of unilateral HS ^35–39^. Epileptiform activity is composed of ictal (seizure) and interictal (between seizures) phases, in which interictal spikes occur (for review see ^40^). In a previous study, we investigated the temporal succession and interaction of interictal and ictal phases in ihKA mice and found that focal seizures, so-called high-load (HL) bursts, formed clusters surrounded by transition phases with short epileptiform (medium (ML) and low-load (LL)) bursts ^41^.

Building on this, we now probe an on-demand LFS protocol that initiates a 10 min stimulation phase as soon as a transition to the ictal state is emerging (evident from an increased interictal spike rate). Here, we provide evidence that on-demand LFS is highly efficient in suppressing focal seizure clusters while strongly reducing the stimulation load compared to continuous stimulation. In addition, we investigated the acute influence of LFS on behavioral performance, including anxiety, spatial navigation, learning, and memory recall. Notably, hippocampal LFS had either positive or neutral effects on hippocampus-related cognitive functions in chronically epileptic mice.

## Material and methods

### Study Design

The objective of this study was to test the therapeutic effects of hippocampal on-demand LFS in a mouse model of epilepsy and compare it to continuous and regular discontinuous LFS stimulation. Male mice of ages 12 to 16 weeks were used for in vivo experiments. Animal procedures were carried out under the guidelines of the European Community’s Council Directive of 22 September 2010 (2010/63/EU) and were approved by the regional council (Regierungspräsidium Freiburg). Mice were chosen randomly for each experimental group (Sal-, Sal+, KA-, KA+). For all experiments, the number of replicates, statistical tests used, and excluded data are reported in the results and figure legends. All LFS protocols were performed sequentially at least twice in each mouse. Owing to the constraints of the experimental design, the experimenter was not blind to manipulations (i.e., electrical stimulations) performed before or during experiments. Analysis, however, was performed in a blinded manner, since only an anonymous identifier for each mouse and trial was given to the different evaluating researchers. No statistical tests were used to predetermine sample sizes of the number of animals, but our sample sizes are similar or larger to those in previous studies.

### Animals

Experiments were conducted with transgenic male mice. For behavioral experiments, we used C57BL/6 Tg(Thy1-eGFP)-M-Line and for LFS protocols Cre-negative littermates from C57BL/6-Tg(Rbp4-cre)KL100Gsat x Ai32(RCL-ChR2(H134R)/EYFP breeding. In total, 106 mice were used for this study. Mice were kept in a 12 h light/dark cycle at room temperature (RT) with food and water ad libitum.

### KA and virus injections

Mice were injected with 50 nl KA (15 mM, Tocris) or saline into the right dorsal hippocampus (LFS protocols: KA n=7; behavioral experiments: KA n=59, saline n=40), as described previously ^35, 37, 39^. In brief, the stereotaxic injection was performed under deep anesthesia (ketamine hydrochloride 100 mg/kg, xylazine 5 mg/kg, atropine 0.1 mg/kg body weight, i.p.) using Nanoject III (Drummond Scientific Company). Mice were randomly assigned to be KA- (15 mM in 0.9% sterile saline) or saline-injected. Stereotaxic coordinates (in mm) were anterioposterior (AP) = −2.0, mediolateral (ML) = −1.5 relative to Bregma, and dorsoventral (DV) = −1.5 relative to the cortical surface. Following KA injection behavioral *status epilepticus* (SE) was verified by observation of mild convulsions, chewing, immobility, or rotations, as described before ^42, 43^. Mice that did not develop SE (n=2), died due to KA treatment (n=27) or surgical procedures (KA n=1, saline n=8), or lost their implant (n=2) were excluded from further experiments.

### Electrode implantations and local field potential recordings

For all experiments, Teflon-coated platinum-iridium wires (125 µm diameter; World Precision Instruments) were implanted 16 days after KA/saline injections into both (ipsilateral and contralateral) dorsal hippocampus (HCi and HCc, respectively) for local field potential (LFP) recordings ^34, 35^. Animals were additionally implanted with a stimulation electrode coated with nanostructured platinum ^44^ directly next to the HCi electrode, but at a 30° angle, targeting the dentate gyrus. Stereotaxic coordinates are given relative to Bregma in mm (AP, ML) or to the cortical surface (DV): AP = −2.0, ML = +1.4 (HCc LFP electrode); -1.4 (HCi LFP electrode); -2.4 (HCi stimulation electrode), DV = −1.6. The correct positions of electrodes and optic fibers were confirmed by post hoc histology (Supplementary Fig. 1). Two stainless steel screws (DIN 84) were implanted above the frontal cortex to provide a reference and ground, respectively. Electrodes and screws were soldered to a micro-connector (BLR1-type). The implants were fixed with dental cement (Paladur).

### Online spike detection and on-demand electrical stimulation

Freely behaving mice were recorded on days 21 and 22 after KA injection (three hours each) to determine reference LFPs. Each mouse represents the biological replicate and the number of recordings per mouse the technical replicate. For LFP recordings, mice were connected to a miniature preamplifier (MPA8i, Smart Ephys/Multi-Channel Systems). Signals were amplified 1000-fold, bandpass-filtered from 1 Hz to 5 kHz, and sampled at 10 kHz (Power1401 analog-to-digital converter, Spike2 software). Subsequently, LFPs were recorded and mice were stimulated on two separate days (three hours each) either discontinuously (10 min 1 Hz “on/off”, week 4 after ihKA injection), on-demand (10 min 1 Hz “on” after initiation, week 5) or continuously (3 h 1 Hz “on”, week 6) at 1 Hz. Three-hour reference LFP recordings were taken at the end of each week. Stimulation experiments consisted of biphasic rectangular current pulses (400 μs phase duration, ± 200 μA final amplitude, anodic first; MC stimulus II software, STG1004, Smart Ephys/Multi-Channel Systems). Each stimulation was started with 25 µA and increased stepwise to 200 µA (25, 50, 100, and 150 µA, 60 pulses each) to avoid the induction of an epileptic seizure (Supplementary Fig. 2).

On-demand stimulation was implemented in custom-made CED-Spike2 and Python scripts running on a desktop PC connected to the data acquisition system (DAQ), similar to ^45^. The Spike2 script downsampled the recorded HCi signal to 500 Hz, transferred the data to the Python script for real-time analysis (adapted from ^41^), and awaited the signal to trigger stimulation (Fig. 1A). Online detection of spikes was performed on 128 pt (256 ms) windows. A new window was evaluated every 64 ms (32 pts). For each detection window, the FFT spectrogram was computed, dynamically normalized to the 5th and 95th reference percentiles per frequency bin, averaged, and then z-scored to the mean and standard deviation (SD) of the average power of the reference. Peaks in the resulting normalized average power exceeding a threshold obtained from the reference were detected as epileptiform spikes. In brief, the reference threshold for spike detection was determined as the center of the plateau region of the spike count vs threshold characteristic (as in ^41^). To account for high amplitude spikes missed by this procedure, deflections larger than 4.5 times the LFP z-scored to the mean and SD of the reference LFP were detected. We set the stimulation threshold to 13 spikes in a 10 s sliding time window. Online detection was paused during the stimulation period and resumed 10 s after the stimulation stop.

**Fig. 1.**
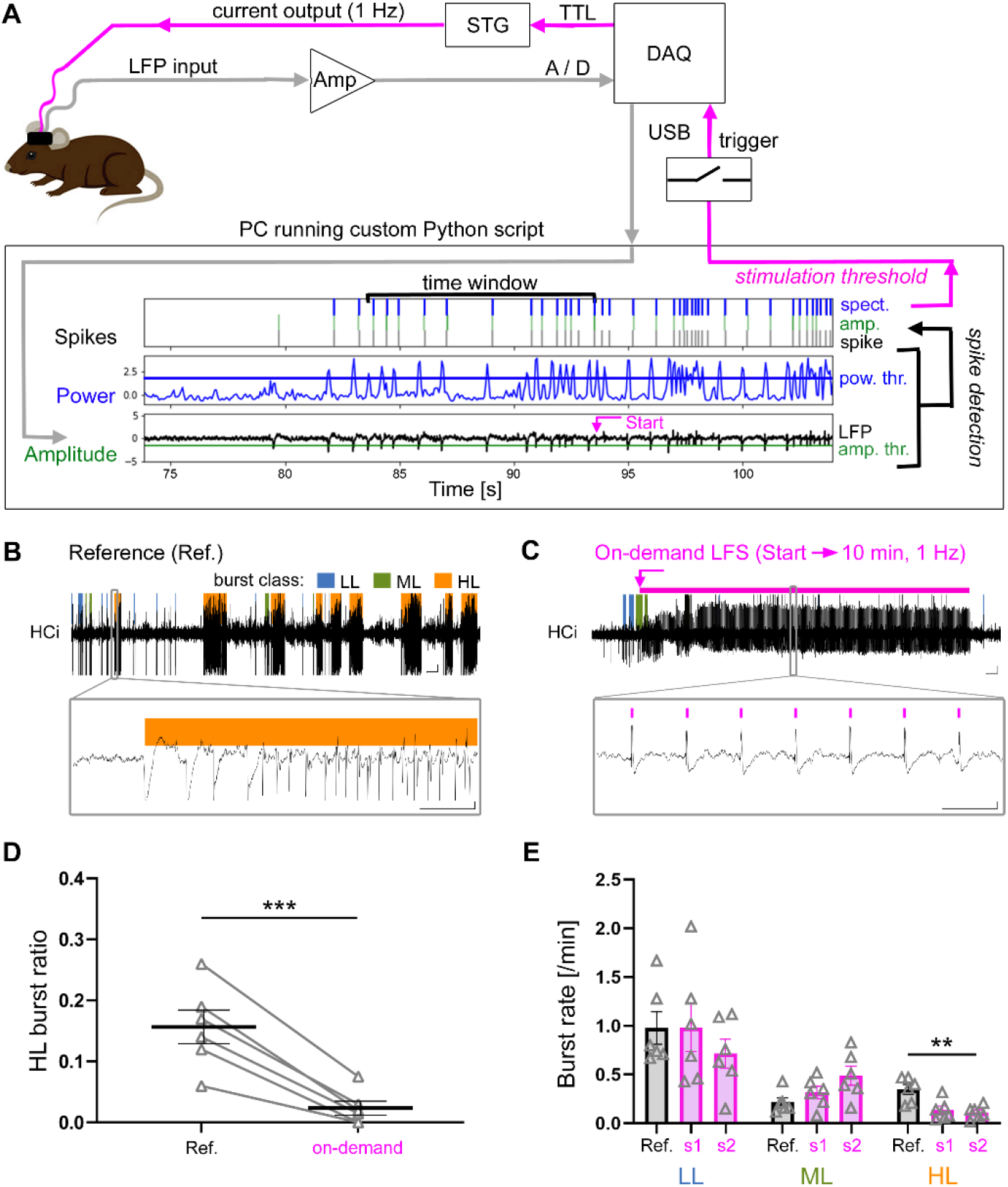
Hippocampal on-demand LFS for the prevention of epileptiform HL clusters. **(A)** Setup design for on-demand stimulation. In brief, 10 min LFS at 1 Hz was initiated if 13 LFP spikes occurred within 10 s. **(B)** Representative LFP trace from an HL (orange) cluster in a reference (Ref.) recording in HCi. Epileptiform activity was categorized offline in LL (blue) ML (green) and HL (orange) bursts. **(C)** Representative LFP trace from an on-demand LFS session with population responses to 1 Hz stimulation (close-up, pink bars). Here, an ML burst initiated LFS (pink arrow: LFS initiation, pink bar: LFS). (B, C) Scale bars: 20 s, 0.5 mV (overview) and 1 s, 0.5 mV (close-up). **(D)** Mean HL burst ratio during reference and on-demand LFS (two three-hour sessions; comparison to three-hour Ref., paired t-test ***p<0.001, n=6 mice, average in black). **(E)** Burst rates of LL, ML, and HLs during Ref. (three hours, gray bar) and on-demand LFS sessions (s1 and s2, three hours, pink bars; multiple t-tests, Holm-Šidák’s correction **p<0.01, n=6 mice). All values are mean ± SEM and individual data points are in grey.

### Behavioral experiments

All behavioral experiments (for timeline see Supplementary Fig. 3). were conducted during the light phase of the day. On experimental days, mice were moved from the animal holding facility to the experimental room at least 30 min before starting the experiments. Mice were handled for five days two minutes per day and reference LFPs were recorded on days 21 and 22 after KA/saline injection (three hours each) as described above. Additionally, we monitored baseline mobility and anxiety levels in the open field on day 21 (15 min, 35x50 cm arena, 60 lux). Animals were video-tracked (Basler acA1300-60gm overhead camera, 60 Hz), with distance, velocity, and time spent in the center (>7 cm away from the arena wall) analyzed with EthoVision XT tracking software (Noldus). The following behavioral tests were preceded by a 30 min LFP recording (Smart Ephys/Multi-Channel Systems 6-channel wireless head stage W2100), during which the stimulated subgroups received continuous 1 Hz LFS (in-house build stimulator) with the parameters described above. Mice were equipped with the wireless head stage during the open field and the light-dark box test.

After four habituation days in the arena (day 25), mice were randomly assigned to be subjected first to either the OLM or the NOR test. The other test was performed after two habituation days on day 38. During training, mice freely explored two identical objects for 10 minutes. After 90 min in the home cage, mice were transferred to the 5-min test trial where one of the objects was moved to a new location or replaced by a novel object. A blinded observer measured the object exploration time (t, defined as the amount of time the mouse’s nose spent touching the object or was <1 cm away from the object). The relative exploration times were expressed as a discrimination index (DI = (t_novel_–t_familiar_)/(t_novel_+t_familiar_)). Objects were cleaned with 0.5% incidin solution (Dr. Schumacher GmbH) after each trial. Trials in which the total exploration time was lower than 2 s (KA- n=1) or mice showed object preference during training (DI>0.3, NOR Sal- n=1, KA- n=1) were excluded from the analysis. Mice that experienced a generalized seizure before the training or the test trial were excluded since those seizures can cause retrograde amnesia by engaging the circuits that participate in memory consolidation ^46^.

On day 27, the light-dark box test was performed to assess the animal’s anxiety-like behavior. Mice were pl0setup and freely explored both chambers (dark: 2 lux and bright: 220 lux, both 18x18 cm) for 10 min. Stimulated subgroups received 30 min LFS directly before the test. The number of transitions between the compartments and the time in light was measured.

Starting on day 29 after KA/saline injections mice were trained in the Barnes maze ^47^. A brightly illuminated (450 lux) open area (platform Ø 100 cm) served as a mild aversive stimulus to encourage the mice to locate an escape box beneath one of the 20 evenly distributed holes along the border of the platform. Eight spatial cues were attached to a black curtain that surrounded the platform. Before the acquisition phase, mice learned to enter the escape box (three times, 2 min habituation). For each trial, mice were placed underneath a start box in the center of the platform for 10 s, the start box was lifted from outside the curtains and the animal had 180 s of exploration time before it was guided to the escape box. The mice then stayed for two minutes in the escape box. Mice were trained three times per day for five consecutive days with an inter-trial interval of 30-40 min. The platform was rotated and cleaned with 0.5% incidin solution between trials. Before the first trial and within the 30 min inter-trial time, mice were LFP recorded and/or stimulated in their home cage. For the test, 24 hours after the last training trial, the target box was removed and mice explored the maze for 90 s. We analyzed the time to target and the number of primary errors. During the test, we additionally measured the time spent in the quadrants of the maze (target, left, right, opposite). The search strategy was defined as either random (random crossings of the platform before visiting the target location), serial (visiting at least two adjacent holes in series before going to the target), or direct (visiting directly or with one adjacent hole next to the target location). Trajectory maps were generated in EthoVision. On days 36 and 37 after KA/saline injection, two three-hour reference LFPs were recorded.

### Tissue preparation and immunohistochemistry

At the end of each experimental series, mice were deeply anesthetized and transcardially perfused with 0.9% saline followed by 4% paraformaldehyde in 0.1 M phosphate buffer (PB, pH 7.4). Following dissection, brains were post-fixated overnight, transferred to PB, and sectioned (coronal plane, 50 μm) on a vibratome (VT100S, Leica Biosystems). Slices were collected and stored in PB for immunohistochemistry.

Free-floating sections were processed for double immunofluorescence staining of neuronal marker (NeuN) and glial fibrillary acidic protein (GFAP). Pre-treatment was done with 0.25% TritonX-100 and 1% bovine serum albumin (Sigma-Aldrich) in PB for one hour. Subsequently, slices were incubated with guinea-pig anti-NeuN (1:1000; Synaptic Systems) and rabbit anti GFAP (1:500; Dako) overnight at 4 °C. On the next day, slices were incubated with anti-guinea pig Cy5 and anti-rabbit-Cy3-conjugated antibodies for 2.5 hours at RT (1:200, Jackson ImmunoResearch Laboratories Inc.) followed by extensive rinsing in PB. Sections were mounted on glass slides with an anti-fading mounting medium (Dako).

### Image acquisition and histological analysis

Tiled fluorescent images of the brain sections were taken with an *AxioImager 2* microscope (Carl Zeiss Microscopy GmbH) using a Plan-Apochromat 10x objective with a numerical aperture of 0.45 (Zeiss). The exposure times (Cy5-labeled NeuN: 800 ms, Cy3-labeled GFAP: 200 ms) were kept constant. In KA mice, the presence of unilateral HS was confirmed in NeuN-labeled sections showing GCD, cell loss in CA1 and CA3, and astrogliosis in GFAP-labeled sections. GCD and astrogliosis were quantified in Fiji ImageJ in three representative sections between -1.58 and -2.06 mm from Bregma. For quantification of the GCL width, three lines perpendicular to the GCL outline were drawn in the upper blade of each NeuN-labeled section. For quantification of astrogliosis, a polygon-shaped region of interest (ROI) was drawn around the pyramidal cell-free gaps in the CA1 region in three representative slices (ipsi and contralateral). Areas with glial scarring around the implantation electrodes were excluded by adjusting the ROI. For normalization, in each slice, the local background was measured in a square (500 pixels^2^) of the contralateral cortex and set as the minimum gray value of each picture. Finally, the integrated density was taken and the mean was calculated for each animal. Animals that showed abnormal contralateral hippocampal atrophy (i.e. extensive loss of dentate granule cells, DGCs) were excluded from the analysis (n=2).

### Offline analysis of epileptiform activity

To detect and classify epileptiform activity in LFPs, we used a semi-automated algorithm that was specifically developed for the ihKA model ^41^. Bursts were classified according to their spike load, hence, three categories of discharge patterns were identified (HL, ML, and LL bursts) as described previously ^35, 41^. To quantify the epileptiform activity within a recording, we calculated the “HL burst ratio” as the fraction of time spent in HL bursts (sum of the HL burst durations divided by total recording time). The automatic detection of HL bursts was confirmed by visual inspection. Generalized seizures and the following 30 min of the LFP recording were excluded from the analysis.

### Statistical analysis

Data were tested for significant differences with Prism 8 software (GraphPad Software Inc.). If data passed the Shapiro-Wilk test for normality (alpha=0.05), comparisons of two groups were performed with a paired (comparisons within animals) or unpaired (comparisons between animals) t-test, otherwise, a Mann-Whitney test was performed. In cases of a comparison of two groups with two subgroups, multiple t-tests with Holm-Šidák correction were performed. If multiple groups were compared, Tukey’s post hoc test was applied. For selected pairs, we chose Šidák’s posthoc test after one-way ANOVA for normally distributed data. If data did not pass the Shapiro-Wilk test, the Kruskal-Wallis test followed by Dunn’s posthoc test was applied. For comparisons of more than two groups over several time points, a two-way ANOVA followed by Tukey’s posthoc test was applied, unless pairs were tested against one selected parameter by Dunnett’s multiple comparisons. Significance thresholds were set to: \**p*<0.05, \*\**p*<0.01, and \*\*\**p*<0.001. For all normally distributed sample populations, the mean and standard error of the mean (SEM) are given, otherwise median with IQR. Correlations were tested using Pearson’s correlation (slope significantly non-zero, confidence interval (CI) 95 %). All statistical results are summarized in Supplementary Table 1.

### Data availability

All data associated with this study are present in the paper or the Supplementary Materials. The raw data are available for research purposes from the corresponding author upon request. The source code files for the offline seizure detection, online spike detection, and on-demand stimulation algorithm are accessible at Zenodo (https://doi.org/10.5281/zenodo.7640783) upon request to K.H.

## Results

### On-demand LFS prevents focal seizure cluster

In this study, we implemented an on-demand 1 Hz electric stimulation of the sclerotic hippocampus intending to suppress clusters of spontaneous focal seizures in chronically epileptic mice. Previous results in the same animal model had shown that the immediate seizure-suppressive effect of LFS was mediated by reduced efficacy of perforant path transmission onto DGCs within the first 10 min of stimulation ^35^. Therefore, we chose a 10 min LFS block to be initiated whenever the threshold of epileptiform spikes was reached (Fig. 1A).

To obtain a reference and calibrate the detection method, we recorded three hours of baseline activity on days 21 and 22, and additional reference LFPs on days 30 and 37 after KA injection (Fig. 1B, Supplementary Fig. 4A,B). Bursts of epileptiform spikes in the ipsilateral hippocampus were automatically classified into LL, ML, and HL bursts. To identify a suitable stimulation threshold for the suppression of focal seizure clusters, we compared the mean epileptic spike rates during the first 10 s of HL and ML bursts of baseline and reference recordings. HL bursts had a much higher spike rate in the first 10 s than ML bursts (Supplementary Fig. 4C, mean ML spike rate 1.11±0.03 Hz; mean HL spike rate 2.77±0.07 Hz, n=6 mice, individual values in Supplementary Table 2). Accordingly, we decided on a threshold of 13 spikes in a 10 s window (corresponding to 1.3 Hz). Between days 33 to 36 after KA injection, we recorded LFPs during three-hour on-demand LFS on two separate days. Before further analysis, stimulation artifacts and stimulation-induced population spikes were detected and excluded (Fig. 1C). To quantify seizure-like activity, we calculated the HL burst ratio, i.e. the total time spent in HL bursts divided by recording length. On-demand LFS strongly reduced the mean HL burst ratio compared to the reference (Fig. 1D, Ref.: 0.16±0.03, on-demand: 0.03±0.01, p<0.001 paired t-test, n=6 mice, mean of two on-demand recordings per mouse). In contrast to a reduction of the HL burst rate, ML and LL burst rates were not affected by on-demand LFS (Fig. 1E, individual values in Supplementary Table 3; multiple paired t-tests with Holm-Šidák correction LL: Ref. vs. on-demand session (s) 1, s2 p=0.986, p=0.363, ML: Ref. vs. on-demand s1, s2 p=0.443, p=0.162 HL: Ref. vs. on-demand s1, s2 p=0.103, p=0.002 n=6 mice).

We observed that a 1 Hz stimulation at 200 µA occasionally induced a focal or even generalized seizure (Supplementary Fig. 2A-D). Ramping up the stimulation current for each LFS start significantly lowered the probability of seizure induction in ihKA-treated mice (Supplementary Fig. 2E, focal seizures: no-ramp 26±8.3%, ramp 0.0±0.0% of started stimulations; F, generalized seizures: no-ramp 25.36±8.25%, ramp 8.0±4.42%, Wilcoxon matched-pairs rank test: both p=0.031, n=10 mice, 3–5 trials per group). In healthy control mice, generalized seizures were induced frequently by 1 Hz at 200 µA but could be completely prevented by the ramp (Supplementary Fig. 2G, no focal seizures with no-ramp and ramp stimuli; H, generalized seizures: no-ramp 70.83±17.18%, ramp 0.0±0.0%, Wilcoxon matched-pairs rank test: p=0.13, n=4 mice, 3–5 trials per group).

Taken together, online detection of epileptiform activity and subsequent initiation of 1 Hz LFS was successful in preventing focal seizure clusters.

### Comparison of on-demand LFS to open-loop LFS

Next, we compared the efficacy of our on-demand LFS protocol in the same mice to open-loop discontinuous (10 min on/off, day 26 to 29) and open-loop continuous (day 40 to 44) 1 Hz-LFS (Fig. 2A). Each corresponding three-hour stimulation protocol was performed twice per week on consecutive days during LFP recordings (Fig. 2B,C). Reference recordings were performed in between LFS protocols to exclude lasting changes in the burst ratio due to LFS (Supplementary Fig. 4D, 21+22d: 0.21±0.04, 30d: 0.16±0.03, 37d: 0.17±0.03, one-way ANOVA F=0.81, n=6 mice, p=0.466). Discontinuous LFS reduced the HL burst ratio only slightly (Fig. 2D, Ref. 21+22d: 0.21±0.04, discontinuous LFS: 0.08±0.03, paired t-test p=0.038) whereas on-demand LFS strongly reduced HL bursts (Fig. 2E, Ref. 30d: 0.16±0.03, on-demand LFS: 0.02±0.01, paired t-test p<0.001). Continuous LFS completely abolished HL bursts (Fig. 2F, Ref. 37d: 0.17±0.03, continuous LFS: 0.00±0.00, paired t-test p=0.003).

**Fig. 2.**
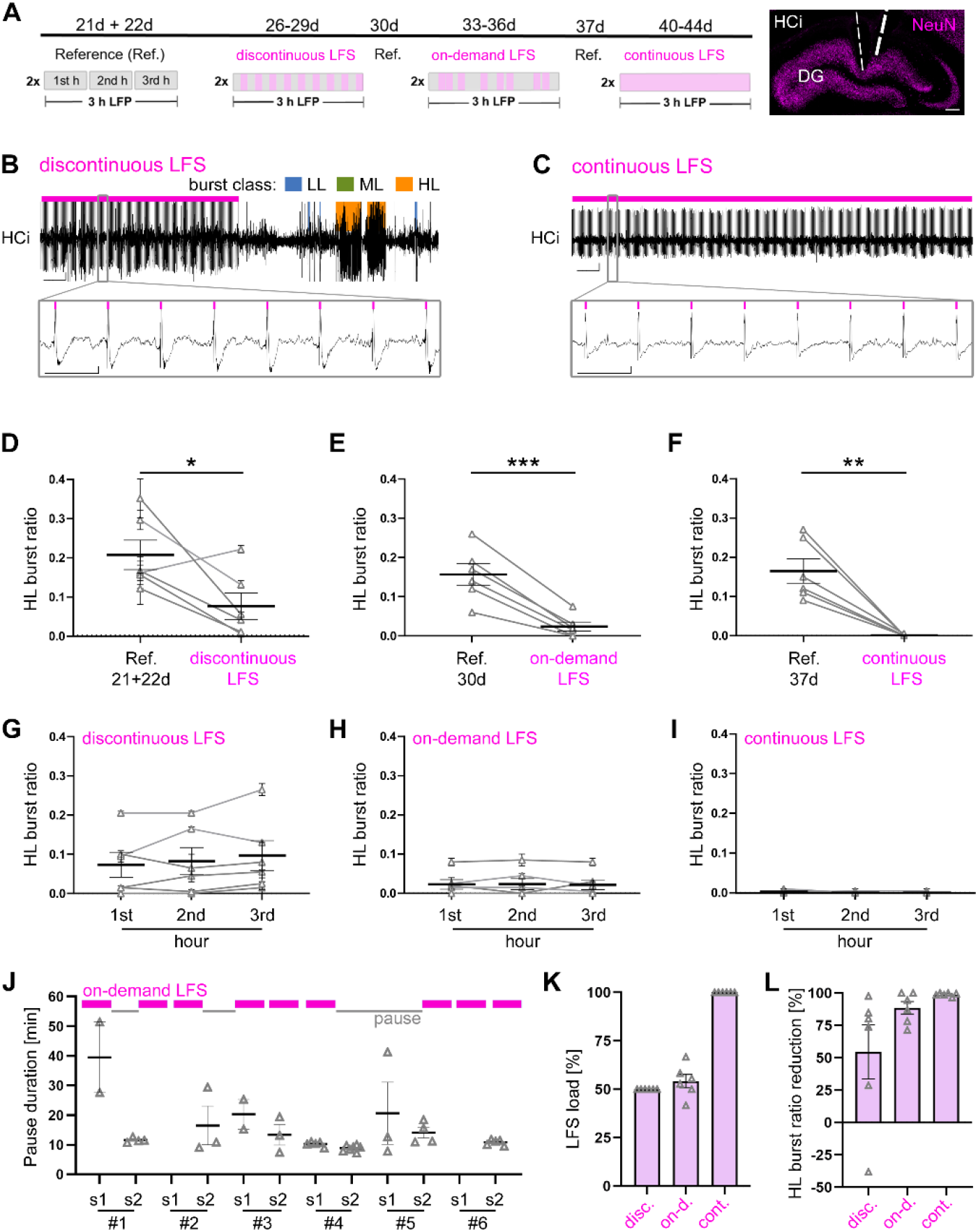
Comparison of on-demand LFS to open-loop discontinuous and continuous LFS. **(A)** Experimental timeline. NeuN staining of an HCi section for validation of HS and electrode positions (LFP left and LFS right). Scale bar: 200 µm. **(B, C)** Representative LFP trace during **(B)** discontinuous and **(C)** continuous LFS in HCi with stimulation artifacts and population responses (pink bars). Scale bars: 20 s, 0.5 mV (overview) and 1 s, 0.5 mV (close-up). **(D)** Mean HL burst ratio of Ref. recordings (two three-hour sessions, day 21 and 22) compared to discontinuous LFS (two three-hour sessions, day 26-29, 10 min on/off; paired t-test *p<0.05, n=6 mice, average in black). **(E, F)** Mean HL burst ratio of Ref. recordings compared to **(E)** on-demand LFS and **(F)** continuous LFS (two three-hour sessions; comparison to three-hour preceding Ref., paired t-test **p<0.01, ***p<0.001, n=6 mice, average in black). **(G, H, I)** Mean HL burst ratio within sessions for **(G)** discontinuous, **(H)** on-demand, and **(I)** continuous LFS. **(J)** Pause durations (median ± interquartile range) during on-demand LFS for individual mice (pauses defined as >5 min). Two sessions were excluded either due to spike detection problems (#2, session 1 (s1) or spontaneous generalized seizure (#6 s1). Inset: Representative session with pink bars for LFS blocks and grey lines for the pauses. **(K)** Percentage of mean LFS time (load) per animal and **(L)** HL burst ratio reduction (n=6 mice). All values are mean ± SEM and individual data points are in grey.

To assess the time course of the stimulation effect, we compared the mean HL burst ratio for the first, second, and third hour of stimulation. Interestingly, the seizure suppressive effect eventuated immediately in all protocols and it did not change over time (Fig. 2G, discontinuous LFS 1^st^ hour: 0.07±0.03, 2^nd^ hour: 0.08±0.04, 3^rd^ hour: 0.10±0.04; one-way ANOVA F=1.7, p=0.24; Fig. 2H, on-demand LFS 1^st^ hour: 0.02±0.01, 2^nd^ hour: 0.02±0.01, 3^rd^ hour: 0.02±0.01; one-way ANOVA F=0.05, p=0.86; Fig. 2I, continuous LFS 1^st^ hour: 0.00±0.00, 2^nd^ hour: 0.00±0.00, 3^rd^ hour: 0.00±0.00; one-way ANOVA F=1.4, all n=6 mice, p=0.3). During reference recordings, burst ratios were constant or increased over time (Supplementary Fig. 4E-G, individual values in Supplementary Table 2).

We analyzed the temporal pattern of LFS blocks and the stimulation load within the three-hour on-demand recordings in all sessions (Supplementary Fig. 5A). LFS was triggered 9.6±0.5 times on average (min.: 6 times; max.: 12 times). The pause durations between LFS blocks showed a bimodal distribution (Supplementary Fig. 5B). 55.81% of all pauses were shorter than 5 min with a mean pause duration of 2.02±0.10 min. This pattern of clustered LFS blocks appeared consistently across individual mice and sessions (Supplementary Fig. 5C). This indicates that half of the time 10 min LFS was not sufficient to reduce epileptiform activity for more than 5 min. Therefore, LFS has been triggered again, elongating the stimulation phases to an overall median LFS time of 20 min (Supplementary Fig. 5D) followed by an extended seizure-free period of 14.25±1.50 min (Fig. 2J). This indicates that the 10 min stimulation blocks in regular 10 min intervals (discontinuous LFS) were too short to reduce epileptiform activity effectively. On-demand LFS, however, had a similar stimulation load than regular, discontinuous LFS (Fig. 2K, discontinuous LFS: 50% of recording time; on-demand LFS: 54.17±3.42% of recording time; continuous LFS: 100% of recording time, n=6 mice) achieving an almost as effective reduction of the HL burst ratio as continuous LFS (Fig. 2L, discontinuous LFS: 54.55±20.88%; on-demand LFS: 88.58±4.82%; continuous LFS: 98,87±0.54%, one-way ANOVA, F=3.94, n=6 mice, p=0.10).

Taken together, while continuous LFS showed the most reliable and reproducible seizure suppressive effect, on-demand LFS was more efficient achieving an 89% burst ratio reduction with only 54% stimulation time.

### Behavioral screening for mobility, anxiety, spatial and non-spatial memory

First, we assessed the mobility and anxiety-like behavior of chronically epileptic and healthy control mice in an open field ^48^. To test whether LFS interferes with hippocampal functions such as learning and memory recall ^49^, we next subjected all mice consecutively to a battery of behavioral tests with and without preceding stimulation, including the light-dark box (anxiety like behavior ^50^), object location (spatial memory ^51^), Barnes maze (spatial navigation and long term memory ^47, 52^) and novel object recognition (non-spatial memory ^53^) (see Supplementary Fig. 3). The stimulation subgroups received 30 min continuous LFS before each training and test session, based on previous observations ^35^, showing that there is at least a 10 min seizure free period after 30 min LFS. Before and at the end of these tests, mice were LFP recorded to acquire the HL burst ratio. After completion of the experimental series, brains were histologically analyzed to quantify HS.

At 21 days after ihKA/saline injection, we video-tracked mice in the open field (Fig. 3A). Healthy control (n=27) and chronically epileptic (n=28) mice were similar with respect to distance (Fig. 3B, saline: 38.37±2.25 m; KA: 35.72±2.55 m; unpaired t-test: p=0.47) and speed (Fig. 3C, saline: 4.55±0.28 cm/s; KA: 4.04±0.29 cm/s; unpaired t-test: p=0.21). Epileptic mice, however, displayed thigmotactic, wall-oriented behavior and spent significantly less time in the center of the field, suggesting an increased anxiety level ^54^ (Fig. 3D, saline: 30.05 [22.16-39.94%]; KA: 12.41 [5.74-21.87%] (median [interquartile range (IQR)]); Mann-Whitney test: p<0.001).

**Fig. 3.**
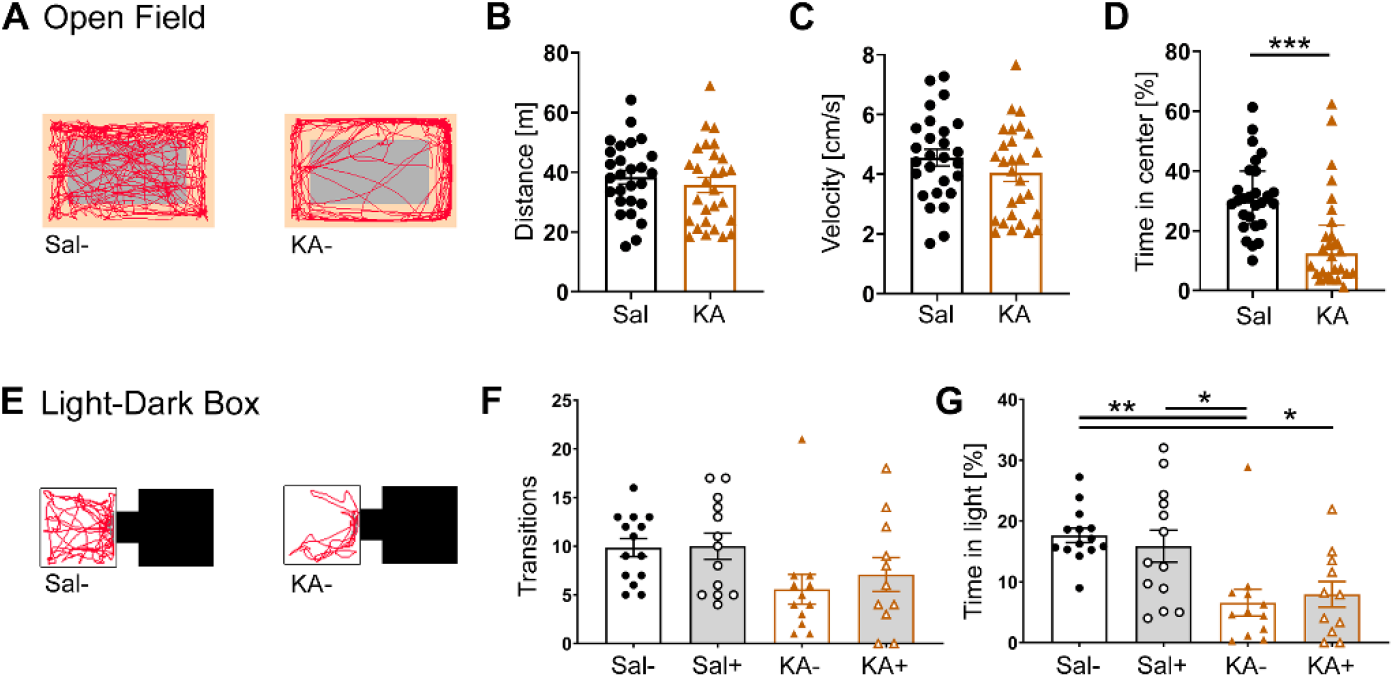
Examination of anxiety-like behavior after hippocampal LFS. **(A)** Representative running tracks (red) in the open field for unstimulated saline (Sal-) and epileptic (KA-) mice 21 days after KA/Sal injection (center area in grey). **(B)** Travel distance, **(C)** velocity, and (D) time in the center region for mice of both groups (Mann Whitney ***p<0.001, Sal-n=27, KA-n=28). **(E)** Representative running tracks (red) during the light-dark box test for Sal- and KA- mice, 27 days after KA/Sal injection. Here, both groups were randomly split into unstimulated (KA-, Sal-) or stimulated (Sal+, KA+) subgroups, the latter receiving continuous hippocampal LFS for 30 min before the test. **(F)** Transitions between the two compartments and **(G)** the total time spent in light for stimulated (Sal+, KA+) or unstimulated (KA-, Sal-) subgroups (one-way ANOVA, Tukey’s multiple comparisons *p<0.05, **p<0.01, Sal- n=14, Sal+ n=13, KA- n=12, KA+ n=11). All values are mean ± SEM, except D shows median ± IQR.

Next, healthy control and epileptic mice were randomly assigned to subgroups with (Sal+ n=13; KA+ n=11) and without stimulation (Sal- n=14; KA- n=12) and were subjected to the light dark box. Mice freely explored the two chambers for 10 min (Fig. 3E). Video tracking revealed that all mice made frequent light-dark transitions (Fig. 3F, Sal-: 9.86±0.93 times, Sal+: 10.0±1.34 times, KA-: 5.58±1.53 times, KA+: 7.09±1.74 times; one-way ANOVA F=2.5, p=0.07). Epileptic mice spent significantly less time in the light, pointing towards elevated anxiety levels compared to healthy controls. In both groups, however, 30 min LFS before the tests did not significantly influence exploration or anxiety-like behavior (Fig. 3G, Sal-: 17.64±1.18%, Sal+: 15.87±2.63%, KA-: 6.58±2.21%, KA+: 7.95±2.11%; one-way ANOVA F=7.24, p<0.001; Tukey’s multiple comparison: Sal- vs. KA- p=0.003, Sal- vs. Sal+ p=0.96, Sal- vs. KA+ p=0.01, KA- vs. KA+ p=0.99, Sal+ vs. KA- p=0.03, Sal+ vs. KA+ p=0.07).

The four experimental groups (Sal- n=14; Sal+ n=13; KA- n=12; KA+ n=11) were next trained in the Barnes maze, a dry-land maze ^52^, adapted for mice ^47^. Mice were trained three times per day for five subsequent days followed by the test session 24 hours later (Fig. 4A). We measured the time to target (primary escape latency) and the number of primary errors (number of holes visited before finding the escape hole for the first time). During the test session, we additionally evaluated the time mice spent in the four quadrants of the maze (target, left, right, opposite) (Fig. 4B).

**Fig. 4.**
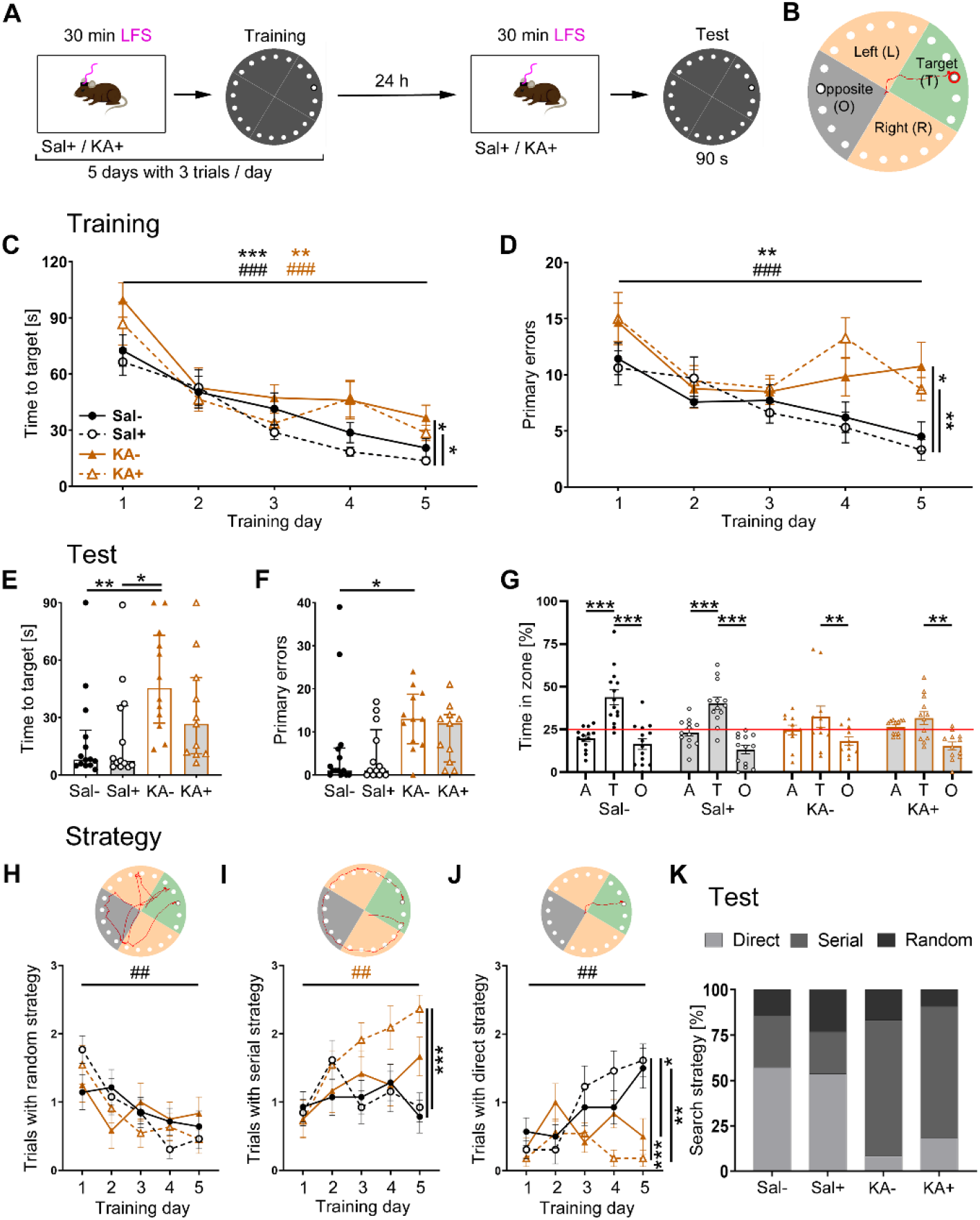
Application of hippocampal LFS prior to spatial learning and memory recall. **(A)** Experimental timeline for the Barnes maze. Sal+ and KA+ mice received 30 min hippocampal LFS before each trial. **(B)** Mice were video-tracked (red trace) on the maze. For the test, 24 hours after the last training trial, the target box (red) was removed. The maze was subdivided into quadrants (target (T), green; opposite (O), grey, and adjacent (A: (left (L)+right (R))/2, orange). **(C)** Time to target and **(D)** primary errors for the experimental groups over five training days (two-way ANOVA comparing training day 1 to day 5 and experimental groups on day 5, Tukey’s multiple comparisons **p<0.01, ***^,###^p<0.001; significance signs: Sal-, black asterisk; Sal+, black hash; KA-, brown asterisk; KA+; brown hash; Sal- n=14, Sal+ n=13, KA- n=12, KA+ n=11 mice). **(E)** Time to target and **(F)** number of primary errors during the test trial (Kruskal-Wallis test, Dunn’s multiple comparisons *p<0.05, **p<0.01, n=C). **(G)** Quadrant preference during the test trial (red line at chance level; two-way ANOVA, Dunnett’s multiple comparisons **p<0.01, ***p<0.001). **(H-J)** The number of trials in which mice used a **(H)** random, **(I)** serial, or **(J)** direct search strategy during training, averaged across mice per day (two-way ANOVA comparing training day 1 to day 5 and experimental groups on day 5, Tukey’s multiple comparisons *p<0.05, **^,##^p<0.01, ***p<0.001, n=C). **(K)** Percentage of mice per experimental group that used random, serial, or direct search strategy in the test trial. All values are given as mean ± SEM, except for H, I, and J, which show individual data points with median ± IQR.

During training, all four experimental groups learned to find the target, since the time to target decreased significantly over five days (Fig. 4C, two-way ANOVA for time and treatment, F(time) (3.02, 138.8)=48.06 p(time)<0.001; Tukey’s multiple comparisons in Supplementary Table 1). In healthy control mice, the number of primary errors decreased continuously with each training day but not in chronically epileptic mice indicating an altered learning behavior (Fig. 4D, two-way ANOVA for errors and treatment, F(errors) (3.32, 152.9)=12.56, p(errors)<0.001; Tukey’s multiple comparisons in Supplementary Table 1). On the fifth training day, stimulated healthy controls were significantly faster at the target than stimulated or unstimulated epileptic animals (Fig. 4C, two-way ANOVA F(treatment) (3, 46)=3.08, p(treatment)=0.04; Tukey’s multiple comparisons in Supplementary Table 1) and made less primary errors (Fig. 4D, two-way ANOVA F(treatment) (3, 46)=4.97, p(treatment)=0.005; Tukey’s multiple comparisons in Supplementary Table 1; individual values for Fig. 4 are reported in Supplementary Table 4). There was no significant influence of LFS on learning behavior in the epileptic or healthy control groups.

Long-term memory performance was examined in the test session, 24 hours after the last training trial (Fig. 4E,F). Chronically epileptic mice showed significantly increased times to target and numbers of primary errors compared to healthy controls indicating deficits in long term memory recall. Interestingly, the performance of stimulated epileptic mice was not significantly different from healthy controls suggesting a positive influence of LFS on memory recall (Fig. 4E, Sal-: 8.06 [5.10-23.40] s, Sal+: 7.36 [4.80-36.16] s, KA-: 45.40 [27.15-72.93]s, KA+: 26.68 [11.12-50.92] s; Kruskal-Wallis test p=0.003; Dunn’s multiple comparisons in Supplementary Table 1; Fig. 4H, Sal-: 1.00 [0.00-6.25] errors, Sal+: 1.00 [0.00-10.50] errors, KA-: 13.00 [7.25-18.75] errors, KA+: 12.00 [3.00-14.00] errors Kruskal-Wallis test p=0.01; Dunn’s multiple comparisons in Supplementary Table 1). We also assessed the preference for the quadrants in absence of the escape box, a marker for the strength of long-term memory. Unstimulated and stimulated healthy control mice stayed significantly longer in the target quadrant than in the adjacent or opposite areas. Unstimulated and stimulated epileptic mice, however, spent significantly less time in the opposite compared to the target zone but not compared to the adjacent quadrants, indicating a weaker memory of the target location (Fig. 4G, two-way ANOVA for time and treatment F(time) (2, 138)=44,20 p(time)<0.001 and F(treatment) (3, 138)=0.26 p(treatment)=0.85; Dunnett’s multiple comparisons in Supplementary Table 1; individual values are reported in Table 2).

Finally, we evaluated the search strategies (random, serial, and direct, see Materials and Methods) during training and test trials. During training, only stimulated healthy control mice increased the number of direct searches from day one to day five (Fig. 4J, two-way ANOVA for searches and treatment F(searches) (3.6,165.4)=5.55, p(searches)<0.001; Tukey’s multiple comparisons in Supplementary Table 1) on the expense of random searches (Fig. 4H, two-way ANOVA for searches and treatment F(searches) (3.46, 159)=9.22, p(searches)<0.001; Tukey’s multiple comparisons in Supplementary Table 1). Stimulated epileptic mice increased the number of serial searches significantly over the five training days (Fig. 4I, two-way ANOVA for searches and treatment F(searches) (3.8, 175)=6.2, p(searches)<0.001; Tukey’s multiple comparisons in Supplementary Table 1). On the fifth training day, the comparison of all experimental groups did not show a main effect regarding the random search strategy (Fig. 4H, two-way ANOVA for searches and treatment F(treatment) (3, 46)=0.14 p(treatment)=0.93). However, stimulated epileptic mice pursued a serial search strategy (Fig. 4I, two-way ANOVA for searches and treatment F(treatment) (3, 46)=2.76, p(treatment)=0.05; Tukey’s multiple comparisons in Supplementary Table 1), whereas healthy controls headed directly to the target (Fig. 4J, two-way ANOVA for searches and treatment F(treatment) (3, 46)=5.03, p(treatment)=0.004; Tukey’s multiple comparisons in Supplementary Table 1). In the test trial (Fig. 4K), chronically epileptic mice used mainly the serial strategy to locate the target (KA- 75.0% and KA+ 72.7%), whereas healthy controls preferred a direct strategy (Sal- 57.1% and Sal+ 53.9%, individual values are reported in Table 2).

Additionally, we assessed spatial learning and short-term memory in the object location test. During a 10-min acquisition phase, mice were exposed to two identical objects. After 90 min in the home cage, mice had 5 min to explore the two objects again, one of which was moved to a novel location (Fig. 5A). Chronically epileptic mice displayed an increased exploration time compared to healthy controls (Fig. 5B, Sal-: 54.61±11.02 s; Sal+: 29.77±8.36 s; KA-: 76.55±9.77 s; KA+: 77.31±14.98 s; one-way ANOVA F=4.7, p=0.01; Tukey’s multiple comparisons in Supplementary Table 1; Sal-, Sal+, KA-, KA+ n=5, 8, 9, 6). None of the groups preferred one object over the other as measured by the discrimination index (Fig. 5C, Sal-: 0.05±0.06; Sal+: -0.05±0.04; KA-: 0.05±0.04; KA+: 0.002±0.03; one-way ANOVA F=1.28, p=0.30). During the test, all experimental groups explored the objects for a similar duration (Fig. 5D, Sal-: 25.48±4.53 s; Sal+: 14.07±1.68 s; KA-: 25.29±3.76 s; KA+: 23.54±2.69 s; one-way ANOVA F=2.03, p=0.13). Only healthy controls clearly discriminated the novel object location (Fig. 5E, Sal-: 0.33±0.05; Sal+: 0.23±0.09; KA-: 0.11±0.10; KA+: 0.19±0.08; one-sample t-test (tested against 0) Sal-: p=0.003; Sal+: p=0.05; KA-: p=0.32; KA+: p=0.08). However, a comparison of the experimental groups showed no differences indicating that preceding LFS did not influence spatial learning and short-term (90 min) memory recall (One way ANOVA F=0.89 p=0.46).

**Fig. 5.**
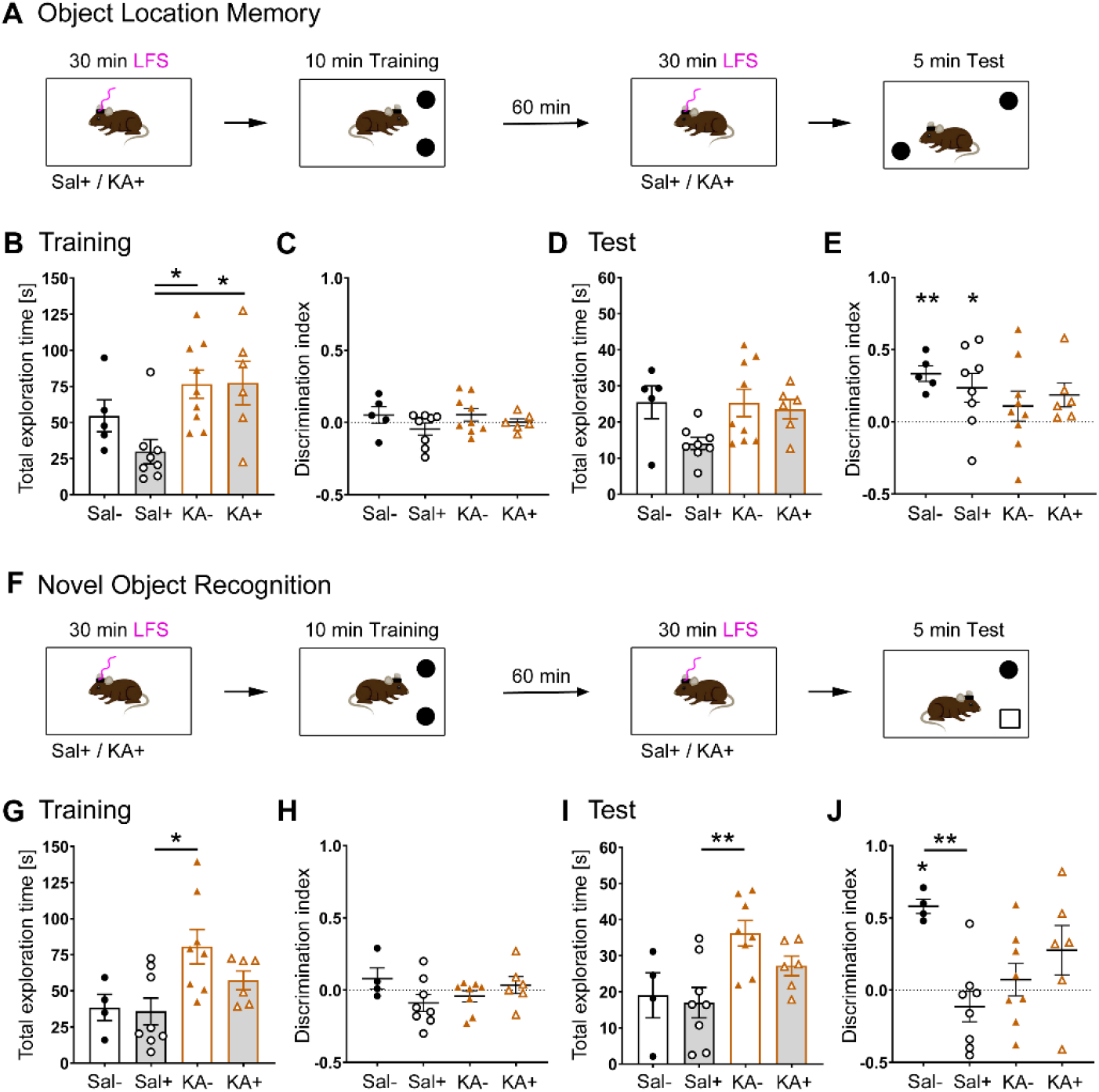
Effects of hippocampal LFS on short-term object location memory and novel object recognition. **(A)** Experimental timeline for object location memory. Stimulated mice (Sal+, KA+) received 30 min hippocampal LFS before each trial. **(B)** Combined exploration time during training (one-way ANOVA, Tukey’s multiple comparisons *p<0.05, Sal- n=5, Sal+ n=8, KA- n=9, KA+ n=6 mice). **(C)** Discrimination index for the training. **(D)** Combined exploration time during the test. **(E)** Discrimination index for the test (one-sample t-test (tested against 0) *p<0.05, **p<0.01). There was no significant difference between the experimental groups. **(F)** Experimental timeline for novel object recognition. One of the two training objects was replaced by a novel object for the test trial. Sal+ and KA+ mice received 30 min LFS before each trial. **(G)** Combined exploration time during training (one-way ANOVA, Tukey’s multiple comparisons *p<0.05, Sal- n=4, Sal+ n=8, KA- n=8, KA+ n=6 mice). **(H)** Discrimination index for the training. **(I)** Combined exploration time during the test (one-way ANOVA, Tukey’s multiple comparisons **p<0.01, n=see G). **(J)** Discrimination index for the test (one-sample t-test (tested against 0) *p<0.05 and one-way ANOVA, Tukey’s multiple comparisons **p<0.01, n=see G). All values are mean ± SEM and individual data points for each mouse.

Finally, we probed non-spatial short-term memory and performed the novel object recognition test (Fig. 5F). During training, epileptic mice explored the objects longer than healthy controls (Fig. 5G, Sal-: 38.55±9.04 s; Sal+: 35.85±9.22 s; KA-: 80.67±11.93 s; KA+: 57.27±6.49 s; one-way ANOVA F=4.56, p=0.01; Tukey’s multiple comparisons in Supplementary Table 1; Sal-, Sal+, KA-, KA+ n=4, 8, 8, 6) but none of the groups displayed object preference (Fig. 5H, Sal-: 0.08±0.07; Sal+: -0.09±0.06; KA-: -0.04±0.04; KA+: 0.04±0.06; one-way ANOVA F=1.67, p=0.20). During the test, unstimulated epileptic mice spent more time exploring the objects than healthy controls (Fig. 5I, Sal-: 19.01±6.22 s; Sal+: 16.98±4.20 s; KA-: 36.21±3.53 s; KA+: 27.19±2.72 s; one-way ANOVA F=5.21, p=0.007; Tukey’s multiple comparisons in Supplementary Table 1). Interestingly, only unstimulated healthy controls could discriminate the novel object (Fig. 5J, Sal-: 0.58±0.05; Sal+: -0.11±0.11; KA-: 0.07±0.11; KA+: 0.28±0.17; one-sample t-test (tested against 0) Sal-: p=0.001; Sal+: p=0.31; KA-: p=0.54; KA+: p=0.17).

Therefore, chronically epileptic mice have impaired short-term object recognition memory. LFS before the training and the test trials caused a slight improvement in chronically epileptic but a strong deterioration in healthy control mice (one-way ANOVA F=4.66, p=0.01; Tukey’s multiple comparisons: Sal- vs. KA- p=0.07, Sal- vs. Sal+ p=0.009, Sal- vs. KA+ p=0.47, KA- vs. KA+ p=0.64, Sal+ vs. KA- p=0.65, Sal+ vs. KA+ p=0.14).

At the end of the behavioral test series, we compared the pathophysiology (HL burst ratio), histopathology (GCD and astrogliosis in CA1), and open-field behavior of stimulated and unstimulated epileptic (KA+ and KA-) mice (Fig. 6) to exclude confounding factors. We found no differences for HS markers (Fig. 6A-F) except a slightly decreased astrogliosis in CA1 of the stimulated group (Fig. 6F). There was no difference in the velocity or time spent in the center among epileptic mice, which were afterward randomly assigned to the LFS subgroups (Fig. 6G,H). Stimulated and unstimulated groups had similar burst ratios in both hippocampi before (Fig. 6I,J) and after (Fig. 6I,K) the behavioral tests (individual values are reported in Supplementary Table 5).

**Fig. 6.**
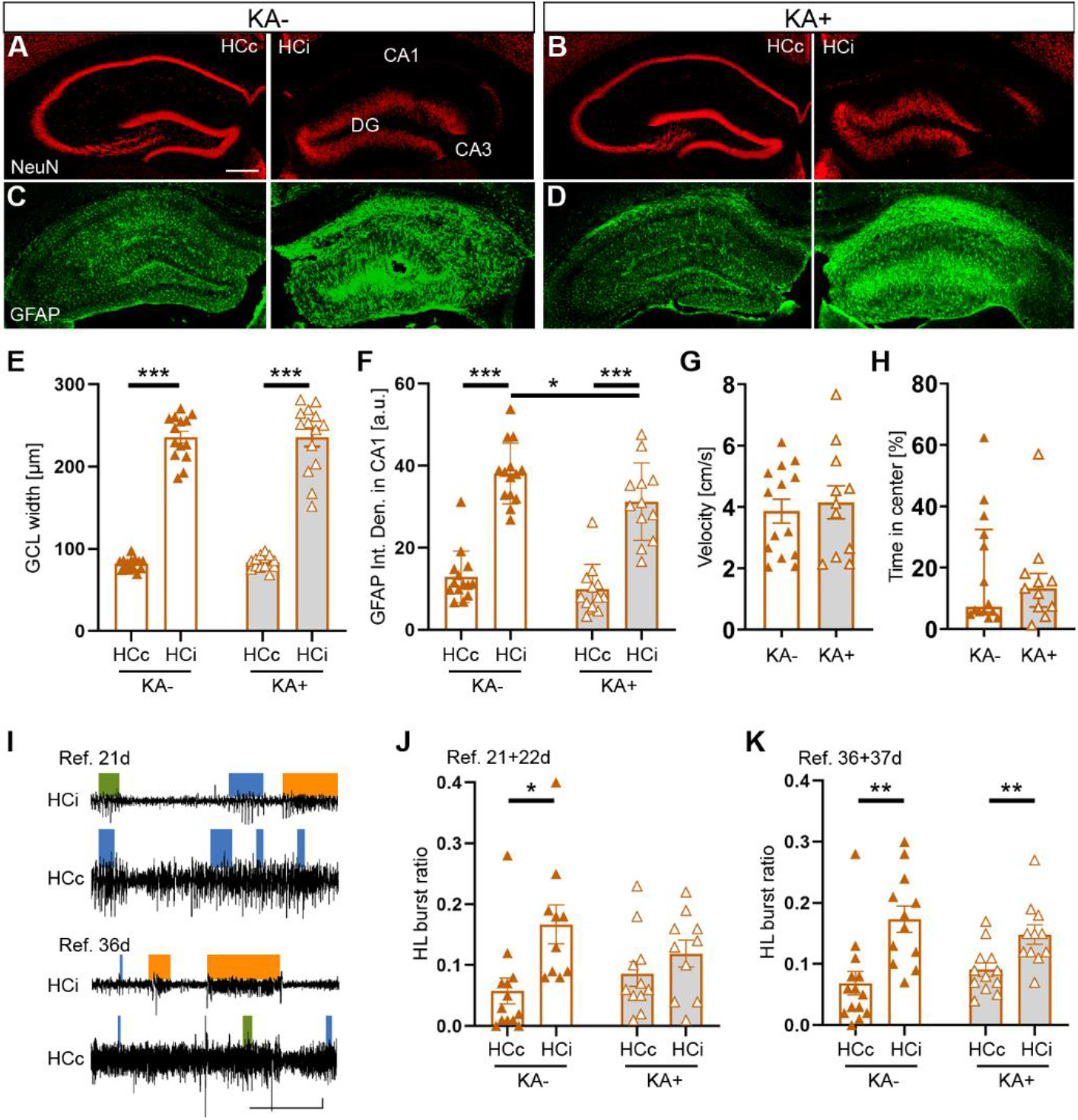
Evaluation of histopathology, pathophysiology, and open-field behavior for epileptic mice subjected to behavioral tests. **(A, B)** NeuN staining visualized cell loss and GCD in HCi while HCc remained unaffected. Scale bar: 200 µm. **(C, D)** GFAP staining for analysis of hippocampal astrogliosis for both, KA- and KA+ mice. **(E)** GCL width for KA- and KA+ mice in HCi and HCc (multiple t-tests, Holm-Šidák’s correction ***p<0.001, KA- n=14, KA+ n=14 mice). **(F)** GFAP integrated density in CA1 for KA- and KA+ mice in both hippocampi (multiple t-tests, Holm-Šidák’s correction ***p<0.001, *p<0.05, KA- n=14, KA+ n=12 mice). **(G)** Velocity and **(H)** time spent in the center area of the open field for mice of both groups (that were later split into KA- and KA+). **(I)** Representative LFP trace from reference (Ref.) recordings in HCi and HCc, 21 days (before LFS and behavioral tests started) and 36 days (after LFS and behavioral tests) post KA injection. Epileptiform activity is categorized into LL (blue), ML (green), and HL (orange) bursts. Scale bar 20 s, 0.5 mV. **(J)** Average HL burst ratio across 6 h of LFP (three hours on 21 and 22 days after KA injection) for KA- and KA+ mice for HCi and HCc (multiple t-tests, Holm-Šidák’s correction *p<0.05, KA- HCc n=13 HCi n=10; KA+ HCc n=11 HCi n=10 mice). **(K)** Average HL burst ratio across 6 h of LFP (three hours on 36 and 37 days after KA injection) (multiple t-tests, Holm-Šidák’s correction **p<0.01, KA- HCc n=14; HCi n=12; KA+ HCc n=12; HCi n=11 mice). All values are mean ± SEM and individual data points for each mouse.

Taken together, in the open field and light-dark box tests chronically epileptic mice showed similar mobility but a higher anxiety-like behavior than healthy controls. In the Barnes maze, they showed an altered navigation strategy and lacked the direct and precise target localization seen in controls. Hippocampal LFS did not aggravate these deficits, indeed, it had a positive influence on long-term memory recall.

## Discussion

The current study explored the therapeutic efficacy of on-demand LFS for seizure suppression and its potential effects on hippocampal function. From a clinical perspective, our findings are of high interest. First, we show that on-demand LFS suppresses clusters of spontaneous, focal seizures as effectively as continuous LFS but with half the stimulation load. Second, we demonstrate that hippocampal LFS does not negatively influence spatial memory formation but

positively affects long-term memory recall in chronically epileptic mice. Therefore, our findings may be useful for the clinical implementation of on-demand LFS in MTLE patients.

### Efficient control of focal seizure clusters by on-demand LFS

A temporally targeted treatment option for MTLE, such as on-demand stimulation, offers several advantages over continuous stimulation since a reduced stimulation time leads to less interruption of normal brain activity and lower power consumption. In clinical practice, RNS^®^ is an FDA-approved closed-loop HFS system that uses power-based measures of LFP activity for seizure detection. It thresholds the line length of the recorded signal, hence it is sensitive to both, frequency and amplitude changes ^55^. However, due to false positive detections, it can apply over 1000 stimulations per day ^17, 56^ which is far beyond the number of seizures patients reported before the start of the therapy ^15^. Successful clinical implementation of on-demand stimulation protocols has three essential requirements: (1) early identification of pathophysiological network patterns in EEG recordings that precede a focal seizure cluster or a generalized seizure, (2) on-demand stimulation of target circuits that prevent their emergence, and (3) exclusion of stimulation-related cognitive impairments.

In the current study, we probed these requirements in a preclinical setting using an MTLE mouse model. In initial tests, we started hippocampal LFS straightaway at a high current (200 µA) which occasionally induced focal or generalized seizures in ihKA mice (25% focal and 25% generalized seizures, Supplementary Fig. 2). This problem has also been observed for DBS in MTLE patients ^26, 57^ and we solved it by ramping up the stimulation current gradually over the first four minutes. To target the first requirement mentioned above, we implemented an online analysis script that identified epileptiform discharges from a single LFP recording electrode using a combination of spectral power and amplitude thresholds. As an alternative to amplitude and/or power-based measures of LFP activity, increased excitability can be measured as a change in phase synchronization ^58–61^. However, this biomarker requires data from multiple recording sites and more computational power. In practice, however, the implantable device is supposed to be used for both, seizure detection and stimulation. Therefore, there are serious constraints for online seizure detection due to the limited availability of computational power and corresponding energy consumption. Here, we focused on detecting interictal epileptiform activity, based on the theory that the transition from an interictal to an ictal state is a slow, dynamic process characterized by the progressive loss of neuronal network resilience ^62^. In ihKA mice, HL bursts of epileptiform activity cluster in time and are surrounded by transition phases consisting of LL and ML bursts ^41^. Concerning long-term dynamics, we observed that the clustering of HL bursts is similar to seizure clusters described for other rodent models ^63^ and patients with drug-resistant focal epilepsy ^58^. Along this line, it was shown that seizures preferentially emerge from distinct brain states and that the effectiveness of curtailing seizures by interneuron stimulation depends on the preceding brain state ^64^. We anticipated that our online detection algorithm identifies ML bursts within a transition phase and starts a 10 min LFS block accordingly. Hence, we initiated LFS during transition phases to stabilize the hippocampal network state at low excitability, thus preventing a focal seizure cluster. In contrast, various studies in other rodent models focused on the identification of seizure onsets with the drawback of delayed action for seizure control ^59–61, 65–68^.

Regarding the second requirement, we applied LFS in the sclerotic hippocampus that, despite the extensive neuronal loss, rewiring, and reactive gliosis, proved to be a suitable target for continuous LFS, suppressing epileptiform activity ^35, 67^. Here, we assessed the efficacy of on demand LFS in comparison to continuous and regular discontinuous LFS. Continuous LFS (100% stimulation) reliably abolished focal seizures by 99%, whereas regular discontinuous LFS (50% stimulation) reduced seizure occurrence by 77%. With on-demand LFS the average total stimulation time was similar to the discontinuous protocol (53% stimulation), while seizure reduction was almost as successful as continuous LFS (-89%) showing that spike rate dependent LFS had a remarkable efficacy for seizure suppression.

Altogether, on-demand LFS presents a highly efficient stimulation protocol based on the detection of an increased interictal spike rate from a single LFP recording electrode in the seizure focus that marks the transition phase to an upcoming focal seizure cluster.

### Hippocampal LFS alleviates deficits in long-term memory recall

Firing patterns and oscillatory dynamics in the entorhinal cortex and hippocampus indicate that this region plays a central role in navigation and memory (for review see ^28, 49^). Therefore, it is crucial to investigate potential behavioral changes that might be associated with hippocampal stimulation used in future therapeutic interventions in MTLE. Human MTLE itself is often associated with behavioral symptoms, such as depression, anxiety, psychosis, and memory impairment ^3, 69^, the severity of which increases with higher seizure frequency and duration of epilepsy ^70, 71^. Anti-epileptic medication or surgical resection for seizure control bears a high risk to affect cognitive performance ^72, 73^. Studies on DBS and RNS^®^ therapies reported variable outcomes on cognition (for review see ^10, 29, 30^) including depression and memory impairment ^74, 75^, which might be related to HFS interfering with ongoing brain activity ^76, 77^. Studies on LFS, reporting stable verbal memory scores with no psychiatric complications, suggest that LFS might be a safer treatment option for pharmacoresistant patients ^22, 23^. However, little is known about whether hippocampal LFS affects navigation and memory.

Here, we used the ihKA mouse model of MTLE that recapitulates the presence of cognitive comorbidities ^78^, especially spatial learning and memory ^79–82^. In our behavioral test series, we first determined the mobility and anxiety levels of chronically epileptic in comparison to healthy control mice. In terms of mobility, epileptic mice were similar to controls but showed elevated anxiety-like behavior in the open field. In the light-dark box, we assessed the influence of LFS on the anxiety level to avoid misinterpretation of cognitive performance ^83^. In fact, LFS did not influence the anxiety level in epileptic and healthy control mice. In the Barnes maze, we observed that healthy controls adapted their search strategy over time to locate the target directly, demonstrating spatial orientation on the maze. In contrast, epileptic mice acquired a different strategy, performing more serial at the expense of direct trials indicating an impairment of spatial navigation ^79^. Possibly, they use local landmarks and self-referential navigation rather than relying on global spatial cues which were also observed in mice with induced mossy fiber sprouting ^84^. In line with studies using the Morris water maze, ihKA-treated mice showed an impairment of long-term (24 hours) memory recall ^78^. In object location memory and novel object recognition tests, studies addressing long-term (24 hours) memory indicated that ihKA mice have problems with locating but not with recognizing objects ^80–82^. In our hands, already short-term (90 min) memory was impaired in both, spatial and non-spatial tests. Therefore, speculating about the underlying mechanism, rewiring of adult DGCs associated with HS ^85^, might impair pattern completion during memory recall ^86^.

Stimulation of the sclerotic hippocampus at 1 Hz before training and test trials did not influence the spatial (object location and target location) and non-spatial (object recognition) learning behavior of epileptic mice. However, LFS impaired non-spatial memory in healthy controls, corroborating that LFS might interrupt mnemonic functions of the dentate gyrus under healthy conditions ^87^. An explanation for this observation might be that LFS mediates inhibition of memory engram formation or recall resulting in false object recognition (for review see ^49^). Importantly, hippocampal LFS alleviated deficits in long-term memory recall (Barnes maze) in chronically epileptic mice. Before learning, mice were stimulated to keep them seizure-free (thus also during learning). Afterward, during memory consolidation, seizures reoccurred and just before memory recall, mice were stimulated again. It remains to be investigated if continuous LFS, e.g. seizure-freedom during the whole consolidation phase, improves their performance or if the lack of CA1 pyramidal neurons and ensuing cortical interactions in the sclerotic hippocampus impairs memory consolidation in general (for review see ^49^).

Our study has some limitations. First, the effects of LFS protocols were probed in defined timescales on focal seizure clusters (three hours) and hippocampal function (30 min). To enhance translatability and safety assessment, future long-term experiments will reveal outcomes of continuously applied LFS protocols on focal as well as generalized seizure reduction and cognitive performance. Second, the implementation of brain state monitoring may improve the detection of increased seizure risk, further improving LFS timing and enabling individual termination. Third, cognitive function includes not only the abilities of learning and memory but also attention, behavioral flexibility, problem-solving, and action planning ^88^, and hence should not be disregarded. A fourth limitation concerns the diurnal variations and sex differences in hippocampal physiology and behavior ^89^, which could potentially influence our findings.

## Acknowledgments

We thank Jessica Link for experimental assistance and data management, Anna Roedling for support with data analysis, and Andrea Djie-Maletz for excellent technical assistance. We are grateful to Prof. Dr. Heinz Beck and André N. Haubrich, University of Bonn, for helpful advice concerning behavioral experiments. We thank Prof. Dr. Maria Asplund, Bioelectronic Microtechnology – IMTEK, for providing laboratory equipment for electrode coatings.

## Supplementary material

Supplementary material is available in a separate file.

Supplementary Fig. 1. Electrode positions for all animals included in the study.

Supplementary Fig. 2. Stepwise increase of stimulation current for the prevention of seizure induction through LFS.

Supplementary Fig. 3. Experimental timeline for behavior experiments. Supplementary Fig. 4. Reference LFPs and spike rates of ML and HL bursts. Supplementary Fig. 5. Temporal pattern of on-demand LFS phases.

Supplementary Table 1. Quantitative summary of statistically tested parameters. (separate excel file)

Supplementary Table 2. Individual values for Supplementary Fig. 4. Supplementary Table 3. Individual values for Fig. 1E. Supplementary Table 4. Individual values for Fig. 4.

Supplementary Table 5. Individual values for Fig. 6.

