## Supplementary material for "On-demand low-frequency stimulation for seizure control: efficacy and behavioral implications"

**List of supplementary Materials:**

Supplementary Fig. 1. Electrode positions for all animals included in the study.

Supplementary Fig. 2. Stepwise increase of stimulation current for the prevention of seizure induction through LFS.

Supplementary Fig. 3. Experimental timeline for behavior experiments.

Supplementary Fig. 4. Reference LFPs and spike rates of ML and HL bursts.

Supplementary Fig. 5. Temporal pattern of on-demand LFS phases.

Supplementary Table 1. Quantitative summary of statistically tested parameters. (separate excel file)

Supplementary Table 2. Individual values for Supplementary Fig. 4.

Supplementary Table 3. Individual values for Fig. 1E.

Supplementary Table 4. Individual values for Fig. 4.

Supplementary Table 5. Individual values for Fig. 6.

### Supplementary Figures

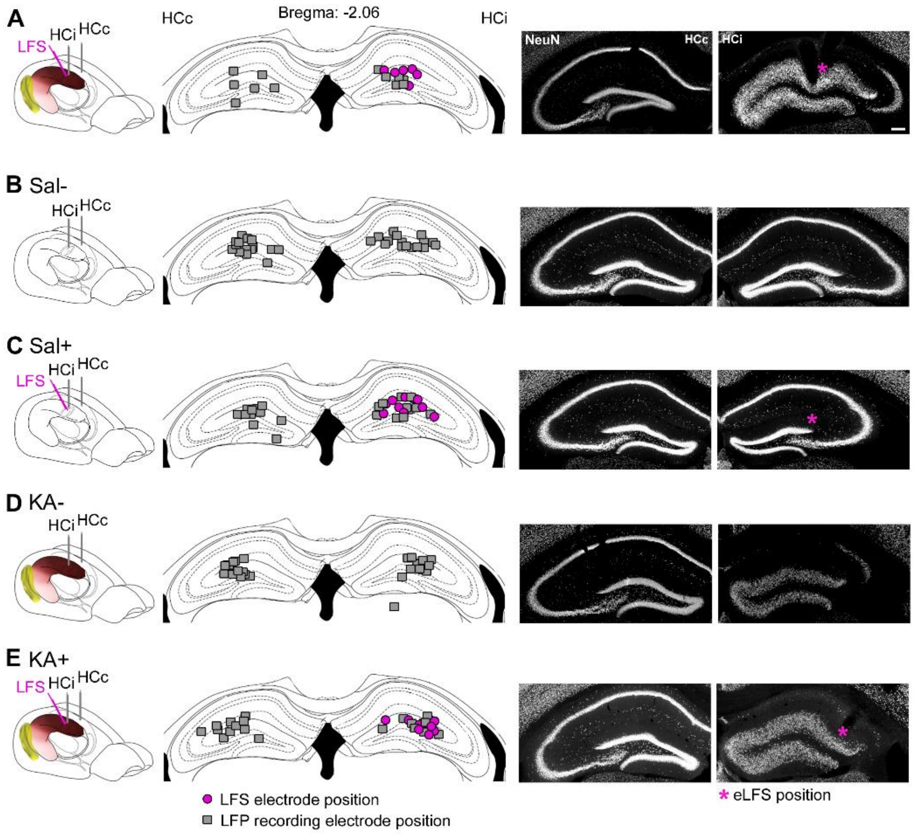


**Supplementary Fig. 1. Electrode positions for all animals included in the study. (A)** LFP (grey) and LFS electrode (pink) positions for all animals included in on-demand LFS experiments. **(B-E)** LFP and LFS electrode positions for all animals included in behavioral experiments. Four experimental groups were used for behavioral experiments: unstimulated (Sal-) and stimulated (Sal+) healthy control mice, as well as unstimulated (KA-) and stimulated (KA+) chronically epileptic mice. Scale bar 200 µm.

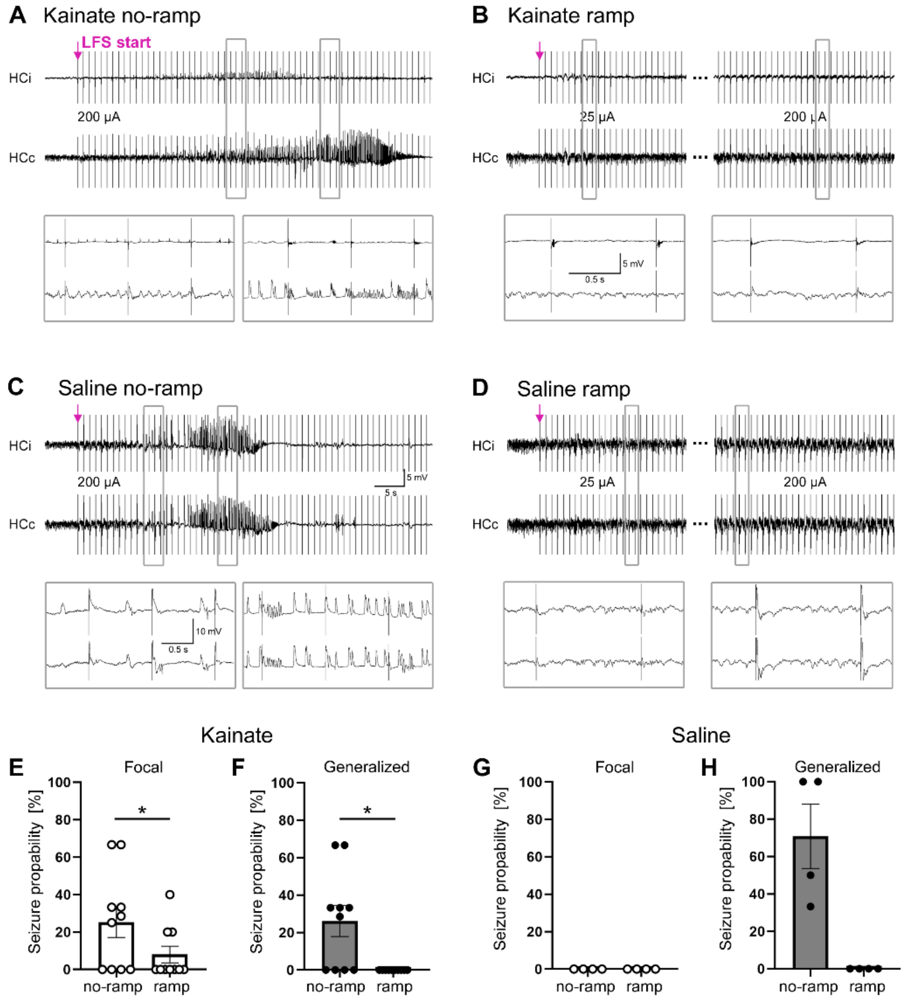


**Supplementary Fig. 2. Stepwise increase of stimulation current for the prevention of seizure induction through LFS. (A, B)** Representative LFP traces from HCi and HCc of a chronically epileptic mouse**. (A)** Starting LFS (1 Hz) at a high current (200 µA, no-ramp) occasionally induced a seizure. **(B)** A stepwise increase of the stimulation current (ramp: 25, 50, 100, and 150 µA, 60 pulses each) prevented seizure induction. **(C)** Representative LFP traces from HCi and HCc of a healthy control mouse that responded with seizure-like activity in HCi and HCc to LFS without the ramp but **(D)** the ramp prevented a seizure induction. **(E, F)** Quantification of seizure probability in KA-treated mice for **(E)** focal and **(F)** generalized seizures for LFS start without a ramp or with a ramp (Wilcoxon matched-pairs rank test, *$p<0.05$, $n=10$). **(G, H)** Quantification of seizure probability in saline-treated mice for **(G)** focal (not occurring) and **(H)** generalized seizures (Wilcoxon matched-pairs rank test, $p=0.13$, $n=4$). Scale bars 5 s, 5 mV (overview) and 0.5 s, 10 mV (close-up). All values are mean ± SEM.

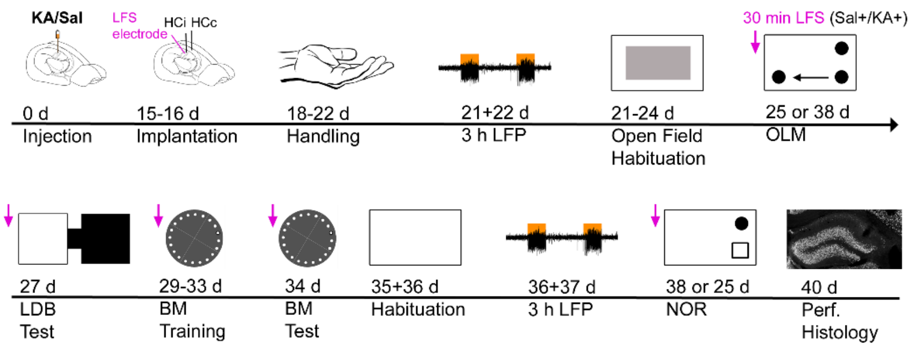


**Supplementary Fig. 3. Experimental timeline for behavioral experiments.** KA- and saline (Sal)-injected mice were equipped with two LFP recording electrodes (HCi and HCc) and one LFS electrode (HCi). Handled mice were LFP-recoded (three hours on days 21 and 22 after KA/Sal injection), tested, and habituated in an open-field arena. Afterward, mice were randomly assigned to the subgroups that were either non-stimulated (Sal-/KA-) or stimulated (Sal+/KA+) with LFS for 30 min before each training and test trial (pink arrow). First, mice were tested in object location memory (OLM) or novel object recognition (NOR), followed by the light-dark box (LDB), Barnes maze (BM), and the missing OLM or NOR test. Mice were randomly assigned to either the OLM or the NOR test first. Two LFP sessions (three hours each) were again recorded on day 36 and 37 days post-KA/Sal injection. After transcardial perfusion, we performed a histological analysis of coronal brain slices.

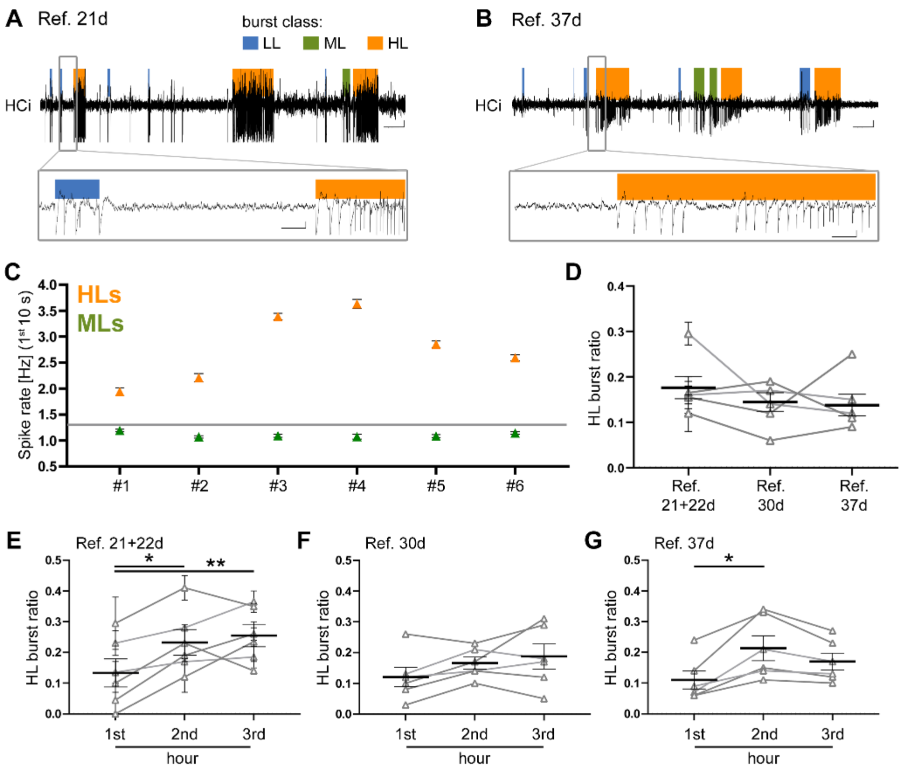


**Supplementary Fig. 4. Reference LFPs and spike rates of ML and HL bursts. (A, B)** Representative reference (Ref.) LFP traces from the ipsilateral hippocampus (HCi) recorded on **(A)** 21 days and **(B)** 37 days after ihKA injection. At both time points, mice are in the chronically epileptic stage, experiencing spontaneous epileptiform activity of all burst classes (LL, blue; ML, green; HL, orange). Scale bars 20 s, 0.5 mV (overview) and 1 s, 0.5 mV (close-up). **(C)** Mean spike rate of the first 10 s from all ML (green) and HL (orange) bursts detected in 12 hours of Ref. recordings (day 21, 22, 30, and 37 after ihKA injection) for each mouse (#1-6). The threshold (grey line, 13 spikes in 10 s) for on-demand LFS was based on these results. **(D)** Mean HL burst ratio for all Ref. recordings. **(E-G)** HL burst ratio for each hour of Ref. recording (one-way ANOVA, Tukey's multiple comparisons *p<0.05, **p<0.01, n=6 mice). All values are mean ± SEM and individual data points for each mouse.

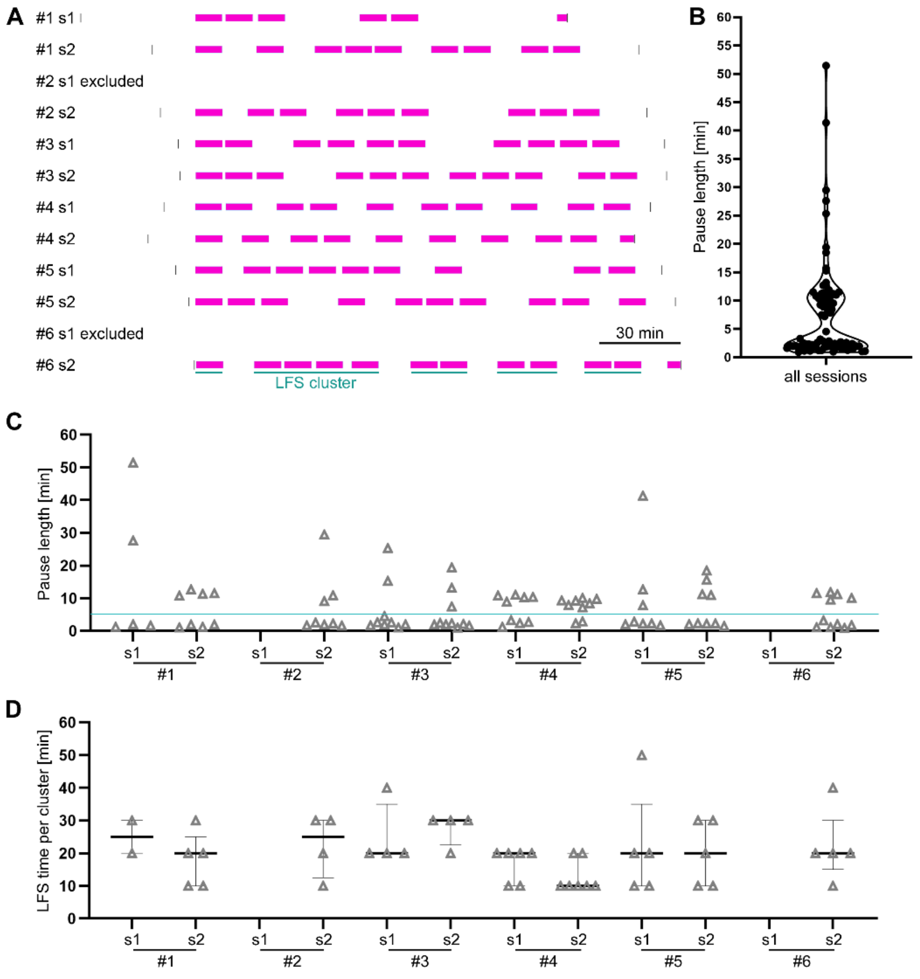


**Supplementary Fig. 5. Temporal pattern of on-demand LFS phases. (A)** Spatial distribution of LFS phases (10 min long, pink bar) for each on-demand session (s1 and s2 for n=6 mice) aligned to the first LFS start. Black bars indicate LFP recording and online detection starts and stops. LFS blocks appeared in clusters (green line). **(B)** Bimodal distribution of pause lengths within all sessions with peaks below and above 5 min. **(C)** Consistent bimodal pattern of pause lengths in each session. **(D)** LFS times per cluster for each session (with median ± IQR) show that LFS was often triggered two or three times in a row, elongating stimulation phases before a pause >5 min occurred.

### Supplementary Tables

##### Supplementary Table 1. Quantitative summary of statistically tested parameters. (separate excel file)

##### Supplementary Table 2. Individual values for Supplementary Fig. 4.

| Fig. S1 C | spike rate | | | | | | |
|---|---|---|---|---|---|---|---|
| ML (1st 10 s) | JL17 | JL18 | JL19 | JL20 | JL21 | JL22 | mean |
| mean [Hz] | 1.19 | 1.07 | 1.09 | 1.07 | 1.08 | 1.14 | 1.11 |
| SEM [Hz] | 0.02 | 0.03 | 0.03 | 0.05 | 0.03 | 0.03 | 0.03 |
| HL (1st 10 s) | spike rate | | | | | | |
| mean [Hz] | 1.94 | 2.21 | 3.39 | 3.63 | 2.85 | 2.59 | 2.77 |
| SEM [Hz] | 0.07 | 0.08 | 0.06 | 0.09 | 0.06 | 0.06 | 0.07 |

| Fig. S1 D | HL burst ratio | | | |
|---|---|---|---|---|
| Refs. | 21+22d | 30d | 37d | n |
| mean | 0.18 | 0.15 | 0.14 | 6 |
| SEM | 0.02 | 0.02 | 0.02 | 6 |

| Fig. S1 E | HL burst ratio | | | |
|---|---|---|---|---|
| 21+22d | 1st hour | 2nd hour | 3rd hour | n |
| mean | 0.13 | 0.23 | 0.26 | 6 |
| SEM | 0.05 | 0.04 | 0.04 | 6 |

| Fig. S1 F | HL burst ratio | | | |
|---|---|---|---|---|
| 30d | 1st hour | 2nd hour | 3rd hour | n |
| mean | 0.12 | 0.17 | 0.19 | 6 |
| SEM | 0.03 | 0.02 | 0.04 | 6 |

| Fig. S1 G | HL burst ratio | | | |
|---|---|---|---|---|
| 37d | 1st hour | 2nd hour | 3rd hour | n |
| mean | 0.11 | 0.21 | 0.17 | 6 |
| SEM | 0.03 | 0.04 | 0.03 | 6 |

##### Supplementary Table 3. Individual values for Fig. 1E.

| Fig. 1 E | Burst rate [/min] | | | | | | | | |
|---|---|---|---|---|---|---|---|---|---|
| | Ref. | | | on-demand s1 | | | on-demand s2 | | |
| | mean | SEM | n | mean | SEM | n | mean | SEM | n |
| LL | 0.98 | 0.17 | 6 | 0.98 | 0.25 | 6 | 0.72 | 0.15 | 6 |
| ML | 0.22 | 0.05 | 6 | 0.32 | 0.06 | 6 | 0.49 | 0.10 | 6 |
| HL | 0.35 | 0.05 | 6 | 0.14 | 0.04 | 6 | 0.11 | 0.03 | 6 |

### Supplementary Table 4. Individual values for Fig. 4.

| Fig. 4 C | Time to target [s] | | | | | | | | | | | |
|---|---|---|---|---|---|---|---|---|---|---|---|---|
| | Sal- | | | Sal+ | | | KA- | | | KA+ | | |
| | mean | SEM | n | mean | SEM | n | mean | SEM | n | mean | SEM | n |
| Day 1 | 72.54 | 8.37 | 14 | 66.42 | 7.18 | 13 | 99.53 | 9.18 | 12 | 86.92 | 11.41 | 11 |
| Day 2 | 50.31 | 8.54 | 14 | 52.77 | 9.46 | 13 | 52.58 | 10.66 | 12 | 46.50 | 6.33 | 11 |
| Day 3 | 41.44 | 8.42 | 14 | 28.83 | 3.91 | 13 | 47.27 | 7.00 | 12 | 33.77 | 5.29 | 11 |
| Day 4 | 28.66 | 5.43 | 14 | 18.48 | 2.36 | 13 | 45.99 | 10.04 | 12 | 46.76 | 9.79 | 11 |
| Day 5 | 20.50 | 4.89 | 14 | 13.63 | 1.84 | 13 | 36.69 | 6.68 | 12 | 28.47 | 4.11 | 11 |

| Fig. 4 D | Primary errors | | | | | | | | | | | |
|---|---|---|---|---|---|---|---|---|---|---|---|---|
| | Sal- | | | Sal+ | | | KA- | | | KA+ | | |
| | mean | SEM | n | mean | SEM | n | mean | SEM | n | mean | SEM | n |
| Day 1 | 11.43 | 1.42 | 14 | 10.62 | 1.53 | 13 | 14.67 | 1.72 | 12 | 15.00 | 2.32 | 11 |
| Day 2 | 7.57 | 0.50 | 14 | 9.69 | 1.89 | 13 | 8.75 | 1.71 | 12 | 9.45 | 1.36 | 11 |
| Day 3 | 7.71 | 1.40 | 14 | 6.62 | 0.91 | 13 | 8.50 | 1.14 | 12 | 8.82 | 1.13 | 11 |
| Day 4 | 6.21 | 1.35 | 14 | 5.31 | 1.37 | 13 | 9.83 | 1.70 | 12 | 13.27 | 1.82 | 11 |
| Day 5 | 4.50 | 1.31 | 14 | 3.31 | 0.92 | 13 | 10.75 | 2.14 | 12 | 8.73 | 1.02 | 11 |

| Fig. 4 G | Zones BM test [%] | | | | | | | | | | | |
|---|---|---|---|---|---|---|---|---|---|---|---|---|
| | Sal- | | | Sal+ | | | KA- | | | KA+ | | |
| | mean | SEM | n | mean | SEM | n | mean | SEM | n | mean | SEM | n |
| Adjacent | 19.85 | 1.72 | 14 | 23.23 | 2.15 | 13 | 26.46 | 1.22 | 12 | 24.66 | 2.89 | 11 |
| Target | 43.85 | 4.37 | 14 | 40.32 | 3.44 | 13 | 31.67 | 3.73 | 12 | 32.44 | 6.24 | 11 |
| Opposite | 16.46 | 3.15 | 14 | 13.22 | 2.46 | 13 | 15.42 | 2.31 | 12 | 18.24 | 2.39 | 11 |

| Fig. 4 H | Random search strategy | | | | | | | | | | | |
|---|---|---|---|---|---|---|---|---|---|---|---|---|
| | Sal- | | | Sal+ | | | KA- | | | KA+ | | |
| | mean | SEM | n | mean | SEM | n | mean | SEM | n | mean | SEM | n |
| Day 1 | 1.14 | 0.25 | 14 | 1.77 | 0.20 | 13 | 1.25 | 0.25 | 12 | 1.55 | 0.28 | 11 |
| Day 2 | 1.21 | 0.26 | 14 | 1.08 | 0.26 | 13 | 0.58 | 0.26 | 12 | 0.91 | 0.21 | 11 |
| Day 3 | 0.86 | 0.18 | 14 | 0.85 | 0.22 | 13 | 1.00 | 0.28 | 12 | 0.55 | 0.21 | 11 |
| Day 4 | 0.71 | 0.19 | 14 | 0.31 | 0.13 | 13 | 0.75 | 0.25 | 12 | 0.64 | 0.20 | 11 |
| Day 5 | 0.64 | 0.23 | 14 | 0.46 | 0.14 | 13 | 0.83 | 0.24 | 12 | 0.45 | 0.21 | 11 |

| Fig. 4 I | Serial search strategy | | | | | | | | | | | |
|---|---|---|---|---|---|---|---|---|---|---|---|---|
| | Sal- | | | Sal+ | | | KA- | | | KA+ | | |
| | mean | SEM | n | mean | SEM | n | mean | SEM | n | mean | SEM | n |
| Day 1 | 0.93 | 0.22 | 14 | 0.85 | 0.19 | 13 | 0.75 | 0.28 | 12 | 0.73 | 0.24 | 11 |
| Day 2 | 1.07 | 0.27 | 14 | 1.62 | 0.29 | 13 | 1.17 | 0.32 | 12 | 1.55 | 0.21 | 11 |
| Day 3 | 1.07 | 0.25 | 14 | 0.92 | 0.24 | 13 | 1.42 | 0.34 | 12 | 1.91 | 0.25 | 11 |
| Day 4 | 1.29 | 0.27 | 14 | 1.15 | 0.30 | 13 | 1.25 | 0.30 | 12 | 2.09 | 0.31 | 11 |
| Day 5 | 0.79 | 0.24 | 14 | 0.92 | 0.21 | 13 | 1.67 | 0.28 | 12 | 2.36 | 0.20 | 11 |

| Fig. 4 J | Direct search strategy | | | | | | | | | | | |
|---|---|---|---|---|---|---|---|---|---|---|---|---|
| | Sal- | | | Sal+ | | | KA- | | | KA+ | | |
| | mean | SEM | n | mean | SEM | n | mean | SEM | n | mean | SEM | n |
| Day 1 | 0.57 | 0.20 | 14 | 0.31 | 0.13 | 13 | 0.33 | 0.14 | 12 | 0.18 | 0.12 | 11 |
| Day 2 | 0.50 | 0.17 | 14 | 0.31 | 0.21 | 13 | 1.00 | 0.28 | 12 | 0.55 | 0.21 | 11 |
| Day 3 | 0.93 | 0.27 | 14 | 1.23 | 0.30 | 13 | 0.42 | 0.15 | 12 | 0.55 | 0.16 | 11 |
| Day 4 | 0.93 | 0.25 | 14 | 1.46 | 0.29 | 13 | 0.83 | 0.21 | 12 | 0.18 | 0.12 | 11 |
| Day 5 | 1.50 | 0.29 | 14 | 1.62 | 0.24 | 13 | 0.50 | 0.26 | 12 | 0.18 | 0.12 | 11 |

| Fig. 4 K | Search strategy BM test | | | |
|---|---|---|---|---|
| | Sal- | Sal+ | KA- | KA+ |
| Direct | 57.14% | 53.85% | 8.33% | 18.18% |
| Serial | 28.57% | 23.08% | 75.00% | 72.73% |
| Random | 14.29% | 23.08% | 16.67% | 9.09% |

**Supplementary Table 5. Individual values for Fig. 6.**

| Fig. S5 E | GCL width [µm] | | | | | |
|---|---|---|---|---|---|---|
| | KA- | | | KA+ | | |
| | mean | SEM | n | mean | SEM | n |
| HCc | 81.78 | 1.92 | 14 | 83.72 | 2.11 | 14 |
| HCi | 235.60 | 7.27 | 14 | 235.48 | 11.00 | 14 |

| Fig. S5 F | GFAP CA1 int. den. [a.u.] | | | | | |
|---|---|---|---|---|---|---|
| | KA- | | | KA+ | | |
| | mean | SEM | n | mean | SEM | n |
| HCc | 12.89 | 1.68 | 14 | 9.92 | 1.75 | 12 |
| HCi | 38.11 | 1.97 | 14 | 31.22 | 2.72 | 12 |

| Fig. S5 G | Velocity [cm/s] | | | | | |
|---|---|---|---|---|---|---|
| | KA- | | | KA+ | | |
| | mean | SEM | n | mean | SEM | n |
| | 3.86 | 0.39 | 14 | 4.15 | 0.54 | 11 |

| Fig. S5 H | Time in center [%] | | | | | |
|---|---|---|---|---|---|---|
| | KA- | | | KA+ | | |
| | median | 25%-75% | n | median | 25%-75% | n |
| | 7.29 | 5.5-32.44 | 14 | 13.25 | 7.18-18.09 | 11 |

| Fig. S5 J | HL burst ratio (21+22d) | | | | | |
|---|---|---|---|---|---|---|
| | KA- | | | KA+ | | |
| | mean | SEM | n | mean | SEM | n |
| HCc | 0.06 | 0.02 | 13 | 0.09 | 0.02 | 11 |
| HCi | 0.17 | 0.03 | 10 | 0.12 | 0.02 | 10 |

| Fig. S5 K | HL burst ratio (36+37d) | | | | | |
|---|---|---|---|---|---|---|
| | KA- | | | KA+ | | |
| | mean | SEM | n | mean | SEM | n |
| HCc | 0.07 | 0.02 | 14 | 0.09 | 0.01 | 12 |
| HCi | 0.17 | 0.02 | 12 | 0.15 | 0.02 | 11 |
